# Towards Digital-Twin-Enabled Bioprocess Monitoring: Fault-Inclusive Soft Sensing of Penicillin Concentration Under Process Deviations

**DOI:** 10.64898/2026.09.21.753176

**Authors:** Abdul Basit Behlim, Atharva Tilewale, Dhaval Patel

**Author notes:** Corresponding Author: Dhaval Patel (,).

## Abstract

Data-driven soft sensors can estimate fermentation product concentrations from routinely recorded process variables, but strong performance during normal operation does not establish reliability during process deviations. This study evaluated current-time penicillin-concentration estimation using 100 simulated IndPenSim batches comprising 90 normal-operation batches and ten documented deviation batches, with 113,935 observations in total. Initial normal-trained models and five follow-up experiments examined complete-batch validation, dependence on batch-progress features, phase-specific error, model-family comparisons, early out-of-distribution warnings and empirical prediction ranges. These analyses motivated a matched comparison between normal-only and fault-inclusive HistGradientBoosting regressors using 36 current and causal history-based inputs. Normal performance was evaluated in five regime-balanced complete-batch folds, and deviation performance by leave-one-fault-batch-out evaluation; every tested deviation batch was excluded from its own model fit. Batch-balanced sample weights were used, with a factor of three assigned to permitted deviation batches in fault-inclusive fitting. On held-out deviation batches, pooled RMSE decreased from 3.195 to 2.564 g/L, a reduction of 19.73%, while MAE decreased from 2.076 to 1.441 g/L and R² increased from 0.8580 to 0.9085. Normal-operation RMSE was nearly unchanged at 1.981 and 1.983 g/L. Eight of ten deviation batches improved. The mean paired fault-batch RMSE difference was −0.7063 g/L, with a descriptive 95% batch-bootstrap interval of −1.2844 to −0.2194 g/L. Improvement was largest within the first-to-last recorded fault-reference window, but late-stage errors persisted. Batch 100 remained poorly predicted, with fault-inclusive RMSE of 6.631 g/L and R² of −2.4837. A fault-risk classifier, Isolation Forest OOD detector and empirical error ranges offered incomplete reliability information. Fault-inclusive training therefore improved this benchmark on average, but did not establish generalization to unseen fault mechanisms, calibrated safety warnings or deployment in physical fermentation.

## 1 Introduction

Fermentation monitoring is most useful when measurements describe both operating conditions and the evolving biological outcome. Temperature, pH, gas composition and flow rates can be recorded frequently, whereas product concentration may be unavailable at the same frequency or only available after laboratory analysis. A soft sensor addresses this gap by estimating a difficult-to-measure variable from signals available at prediction time. Reviews of process and bioprocess soft sensing emphasize that usefulness depends not only on average accuracy but also on data quality, maintenance, changing batch phases and reliability under conditions that differ from model development (Brunner et al., 2021; Kadlec et al., 2009; Luttmann et al., 2012).

Penicillin concentration is a natural target because it represents product accumulation through a long fed-batch trajectory. A model can nevertheless appear accurate by learning typical progress patterns and then continue to predict normal production after a deviation has suppressed the simulated product response. IndPenSim provides a reproducible setting for studying this problem. The simulator represents an industrial-scale fed-batch penicillin process, and the public benchmark contains three normal operating strategies and documented process-deviation batches (Goldrick, 2019; Goldrick et al., 2019; Goldrick et al., 2015). These data are simulated; they are not a substitute for physical-plant validation.

Prior penicillin soft-sensor research includes feature-selected recurrent models, multi-output support-vector frameworks, forecasting approaches, MLOps-based maintenance and interpretable tree learners (Acosta-Pavas et al., 2024; Hua et al., 2023; Li et al., 2023; Metcalfe et al., 2025; Rekkas-Ventiris & Benardos, 2026). This literature establishes that penicillin concentration modelling and abnormal-operation monitoring are not new. The present question is narrower: when the model family, features and outer validation unit are held fixed, does including permitted deviation batches during training improve current-concentration estimates on a completely held-out deviation batch without materially degrading held-out normal batches?

Hua et al. combined Random Forest feature selection with an optimized long short-term memory network for penicillin fermentation (Hua et al., 2023). Li et al. developed a multi-output representation-learning and support-vector-regression framework (Li et al., 2023), while Rekkas-Ventiris and Benardos studied time-series forecasting of penicillin concentration (Rekkas-Ventiris & Benardos, 2026). Those studies use different prediction formulations and should not be treated as matched competitors unless reimplemented with the same inputs, target timing, units and complete-batch partitions. Within IndPenSim, Metcalfe et al. embedded an LSTM soft sensor in an MLOps workflow and used batches 91–100 to examine deterioration and maintenance (Metcalfe et al., 2025). Acosta-Pavas et al. compared interpretable learners, including tree-based methods, on industrial-scale fermentation simulations (Acosta-Pavas et al., 2024). The current study complements these analyses by using the same HGB settings for the two principal training strategies and pairing the resulting errors at the held-out-batch level. It retains unsuccessful cases rather than reporting only pooled improvement.

Siegl et al. developed an adaptive ensemble soft sensor for biomass monitoring in Pichia pastoris, with reliability-based handling of sensor faults (Siegl et al., 2022). Jin et al. proposed online ensemble Gaussian-process regression for time-varying batch processes and demonstrated adaptation in an industrial fed-batch chlortetracycline setting (Jin et al., 2015). These are important precedents for fault tolerance and model adaptation, but they address changing models or sensor reliability rather than the present offline comparison of normal-only and fault-inclusive concentration regressors. Rivera et al. combined process analytical technology signals with a neural soft sensor for intensified sugarcane-ethanol fermentation (Rivera et al., 2024). Del Hierro et al. evaluated real-time biomass estimation across 19 independent high-density yeast batches using a nested leave-one-batch-out design (Del Hierro et al., 2026). These physical-fermentation studies show the value of batch-level validation and online variables in settings beyond simulation. Their experimental systems, targets and sample sizes differ, so they provide methodological context rather than direct numerical benchmarks. Liu et al. proposed an LSTM-based fault-tolerant soft sensor for uneven batch processes under sensor failure (Liu et al., 2024). Sensor failure, process deviation and distribution shift can all harm prediction, but they are not interchangeable. Kadlec et al. describe adaptation mechanisms for data-driven soft sensors (Kadlec et al., 2011), whereas process-history fault-diagnosis literature focuses on detecting and diagnosing abnormal states (Venkatasubramanian et al., 2003). In the present work, the HGB regressor estimates concentration; the separate warning score and OOD detector do not identify physical root causes.

Repeated observations within a fermentation batch are serially related and do not constitute independent production runs. Structured cross-validation guidance therefore supports keeping complete groups outside fitting (Roberts et al., 2017). Leakage can otherwise create optimistic machine-learning estimates, particularly when preprocessing or selection decisions use the evaluation data (Kapoor & Narayanan, 2023). The batch is consequently the outer validation unit here, and imputation is fitted on each training partition. Random Forest, gradient boosting and Isolation Forest are established algorithms (Breiman, 2001; Friedman, 2001; Liu et al., 2008), implemented with scikit-learn (Pedregosa et al., 2011). No new learning algorithm is claimed. A complete-batch bootstrap is used only as a descriptive summary of paired batch errors (Efron, 1979). Reliability terminology is similarly restricted: empirical residual ranges are not a decomposition of aleatoric and epistemic uncertainty (Hüllermeier & Waegeman, 2021), dataset shift can undermine uncertainty estimates (Ovadia et al., 2019), and an OOD score marks unusual inputs rather than a confirmed process fault (Yang et al., 2024). The classifier output is explicitly treated as uncalibrated because numerical scores do not become probabilities without calibration assessment (Guo et al., 2017). ROC AUC is reported within held-out fault batches, but precision-recall analysis would be important for deployment with imbalanced warnings and is not reported here (Saito & Rehmsmeier, 2015).

Relevant precedents also exist outside IndPenSim. Adaptive ensemble soft sensors have been evaluated for fault-tolerant biomass monitoring, online ensemble Gaussian-process regression has been used for nonlinear time-varying batch processes, and soft sensors have been studied with experimental ethanol and high-density yeast fermentations (Del Hierro et al., 2026; Jin et al., 2015; Rivera et al., 2024; Siegl et al., 2022). A separate LSTM study addressed sensor-failure tolerance in uneven batch processes (Liu et al., 2024). Reviews distinguish such adaptation or fault diagnosis from the simpler act of adding abnormal examples to an offline regression training set (Kadlec et al., 2011; Venkatasubramanian et al., 2003). These studies motivate robust evaluation, but their targets, sensors, units, fault definitions and validation designs differ from the current benchmark, so their reported errors cannot be ranked directly against the present RMSE values.

The digital twin is now considered as a promising framework to combine process models, routine measurements and data-driven estimators to describe the evolving bioprocess. In this context, soft sensors can be used to add an additional layer of state-estimation, by inferring biological or product target variables that are not easily measurable, from signals that are standardly accessible. However, a soft sensor designed to operate optimally during nominal operation can give erroneous state estimates when the process enters an operation region that is not covered by the model training data set. Such robustness to parameters or operation transitions faced during production is thus relevant for digital-twin-based monitoring, especially when the estimated states will then be used for forecast, decision or control.

For instance, IndPenSim provides a reproducible dynamic fermentation environment in which processing trajectories, normal operating strategies and documented deviations can be analyzed. We used this environment to test whether adding allowed deviation trajectories in the soft-sensor training set improves the current-time penicillin-concentrations estimation on held-out deviation batches, without losing accuracy on normal operation. The present study addresses this specific digital-twin-enabling component rather than implementing a complete digital twin.

The work proceeded in two stages. First, a normal-trained baseline and five diagnostic experiments examined whether the observed weakness could be explained by one split, two progress variables or one model family, and whether early OOD and empirical intervals were informative. Second, the main analysis compared normal-only and fault-inclusive HGB within a complete-batch outer evaluation. The same finite benchmark informed both stages, so the work is an exploratory benchmark study rather than an external confirmation. Three questions were addressed. First, does fault-inclusive, batch-weighted fitting reduce deviation-batch concentration error without a material loss on normal batches? Second, are improvements shared across batches and fault phases or dominated by a few trajectories? Third, do auxiliary warning scores identify unreliable predictions? The task is current-time state estimation rather than forecasting or automatic root-cause diagnosis. Accordingly, the study evaluates a soft-sensing and reliability-monitoring component relevant to a future bioprocess digital twin, rather than claiming implementation or validation of a complete digital-twin system.

## 3 Materials and methods

### 3.1 Dataset and target

The public 100-batch IndPenSim benchmark was analyzed (Goldrick, 2019). Batches 1–30 use recipe-driven control, batches 31–60 operator-controlled operation and batches 61–90 advanced process control. Batches 91–100 form the documented deviation group. The retained process exports contained 113,935 observations sampled every 0.2 h, equivalent to 12 min. Last recorded time ranged from 167 to 290 h. These rows are repeated measurements from 100 simulated trajectories, not 113,935 independent experiments.

**Table 1:** Dataset groups used in the analysis.

| Operating group | Batch IDs | Batches | Observations |
| --- | --- | --- | --- |
| Recipe driven | 1–30 | 30 | 34,325 |
| Operator controlled | 31–60 | 30 | 34,385 |
| Advanced process control | 61–90 | 30 | 33,700 |
| Documented deviation | 91–100 | 10 | 11,525 |
| Total | 1–100 | 100 | 113,935 |

*Batch design follows the public IndPenSim benchmark. Counts were calculated from the retained process exports. All batches are simulated.*

The target was the same-time dataset variable Penicillin concentration in g/L. The original export contained 39 process or metadata columns and 2,200 Raman columns. Raman spectra were excluded to define a process-variable soft sensor. The working split exports contained 41 columns after administrative batch and split labels were added; those labels and all fault-reference fields were excluded from regression inputs.

### 3.2 Batch identification splitting and preprocessing

Batch identities were reconstructed from resets in elapsed time and checked against the expected sequence 1–100. Observations were sorted by batch and time before any history variables were generated. The original development split used 60 normal training batches, 15 normal validation batches, 15 normal test batches and ten deviation stress-test batches, containing 67,820, 17,510, 17,080 and 11,525 rows respectively. Regimes were balanced across the three normal subsets and no batch overlapped between them. Exact memberships are reported in Supplementary Table S2.

Non-finite inputs were converted to missing values. Median imputation was fitted on the training part of every reported model fit and then applied to held-out batches. Structural missing values at batch starts were retained until imputation; no backward filling from later observations was used. Batch identity, regime, split label, fault reference and fault flag were not predictors. Feature-usability screening based on missingness and variation was performed on the original baseline training partition, and the retained list was reused across the 25 repeated Random Forest fits rather than re-established inside each fold. Those repeated results therefore condition on a fixed feature set and do not validate the complete screening procedure.

The documented fault-batch designation was taken from the benchmark design. An earlier binary rule based on the Fault flag column had incorrectly marked all batches as faulty; it was corrected before the reported split files were created. Within batches 91–100, non-zero Fault reference observations were used only for retrospective phase labels. Missing reference values were treated as zero, and the reference was never supplied as a model input.

### 3.3 Current and history-based inputs

The final regressor used 36 inputs: 20 current or base variables, 15 history variables and one cumulative-feed variable. Five signals—dissolved oxygen, sugar-feed rate, temperature, pH and aeration—each contributed a one-step lag, a first difference and the mean of up to five preceding observations. The previous-five mean excluded the current row by shifting once before the rolling calculation. At the 0.2 h sampling interval, a full five-observation history spans the preceding hour. All transformations were grouped within batch, and no lagged penicillin target was used.

**Table 2:** Input feature families for the final regressor.

| Feature family | Count | Information used |
| --- | --- | --- |
| Current or base inputs | 20 | Current process values and elapsed time |
| One step lags | 5 | Previous observation of each selected history signal |
| First differences | 5 | Current minus previous signal value |
| Previous five means | 5 | Up to five preceding signal values |
| Cumulative feed | 1 | Within batch feed rate multiplied by positive time increments |
| **Total** | **36** | **Current concentration inputs** |

Cumulative sugar feed was computed by multiplying the current volumetric sugar-feed rate by the positive elapsed-time increment and cumulatively summing within each batch; the first increment was zero. Despite its saved name, its resulting unit is liters of feed, not grams of sugar, because no feed-concentration conversion was applied. Both elapsed time and cumulative feed can encode progress through a batch. Their value was examined by ablation, but their importance does not establish a biological causal mechanism.

### 3.4 Initial model development

A training-median dummy regressor, multiple Linear Regression and Random Forest were compared on the 15-batch normal validation subset. Linear Regression used training-fitted imputation and standardization. Random Forest used median imputation, 150 trees, maximum depth 18, minimum leaf size two and random seed 42. The model with the lowest validation RMSE was selected and refitted on the combined 75 normal training and validation batches. The selected model was then evaluated on the fixed 15-batch normal test set and ten-batch deviation stress test. Only the selected Random Forest underwent detailed testing in the initial section; a later experiment generated final test results for all four comparator families.

### 3.5 Five diagnostic experiments

Experiment 1 repeated regime-stratified complete-batch validation for Random Forest. Within each of five repetitions, batches in each normal regime were independently shuffled using seeds 42–46 and divided into five groups of six. One group from each regime formed an 18-batch test fold, leaving 72 normal batches for training. This produced 25 fits, with every normal batch tested once per repetition. The 25 estimates are dependent because they reuse the same 90 trajectories.

Experiment 2 refitted Random Forest with all 36 inputs, without time, without cumulative feed and without both. All variants used the same 75 normal development batches and fixed normal and deviation test batches. The experiment examines two explicit progress features but does not remove every possible progress proxy. Experiment 3 divided each documented deviation trajectory into rows before the first active fault reference, the inclusive first-to-last activity window, and rows after the last activity. Inactive gaps between episodes remained inside the window. The phase labels are retrospective and do not estimate a causal fault effect.

Experiment 4 compared the median dummy, Linear Regression, Random Forest and an earlier normal-trained HGB implementation. That HGB used learning rate of 0.05, 300 boosting iterations, 31 maximum leaves, minimum leaf size 20, L2 regularization 1 and no early stopping. Experiment 5 fitted separate early-batch OOD detectors at 12, 24, 48 and 72 h. For each horizon and batch, the mean, standard deviation and last observed value of 35 inputs yielded 105 summary features. Training-median imputation, standardization, PCA retaining at least 95% variance and a 500-tree Isolation Forest were fitted using the 75 normal development batches. The threshold was the fitted reference 95th percentile. A separate Random Forest empirical range used 60 normal training batches and 15 normal calibration batches with nominal 90% coverage.

### 3.6 Main fault-inclusive HGB comparison

The main analysis used scikit-learn HistGradientBoostingRegressor for both strategies. Each model used squared-error loss, learning rate 0.05, 180 boosting iterations, 31 maximum leaves, minimum leaf size 30, L2 regularization 1 and no early stopping. Hyperparameters were fixed before outer evaluation. The normal-only and fault-inclusive regressors received identical feature definitions and estimator settings; only the permitted training observations and weights differed.

Within each fit, a row in batch b received a base weight proportional to 1 divided by the number of rows in that batch, so every batch contributed the same total base mass. The multiplier m(b) was one for a normal batch and three for a deviation batch in fault-inclusive fitting; weights were then normalized to mean one. In a leave-one-deviation-batch-out fit, the 90 normal batches contributed relative mass 90 and the nine permitted deviation batches mass 27, corresponding to 23.08% of total weight. The multiplier was not tuned in an outer loop. Because fault inclusion and up-weighting changed together, the analysis cannot separate the effect of seeing deviation examples from the effect of assigning them additional weight.

**Table 3:**
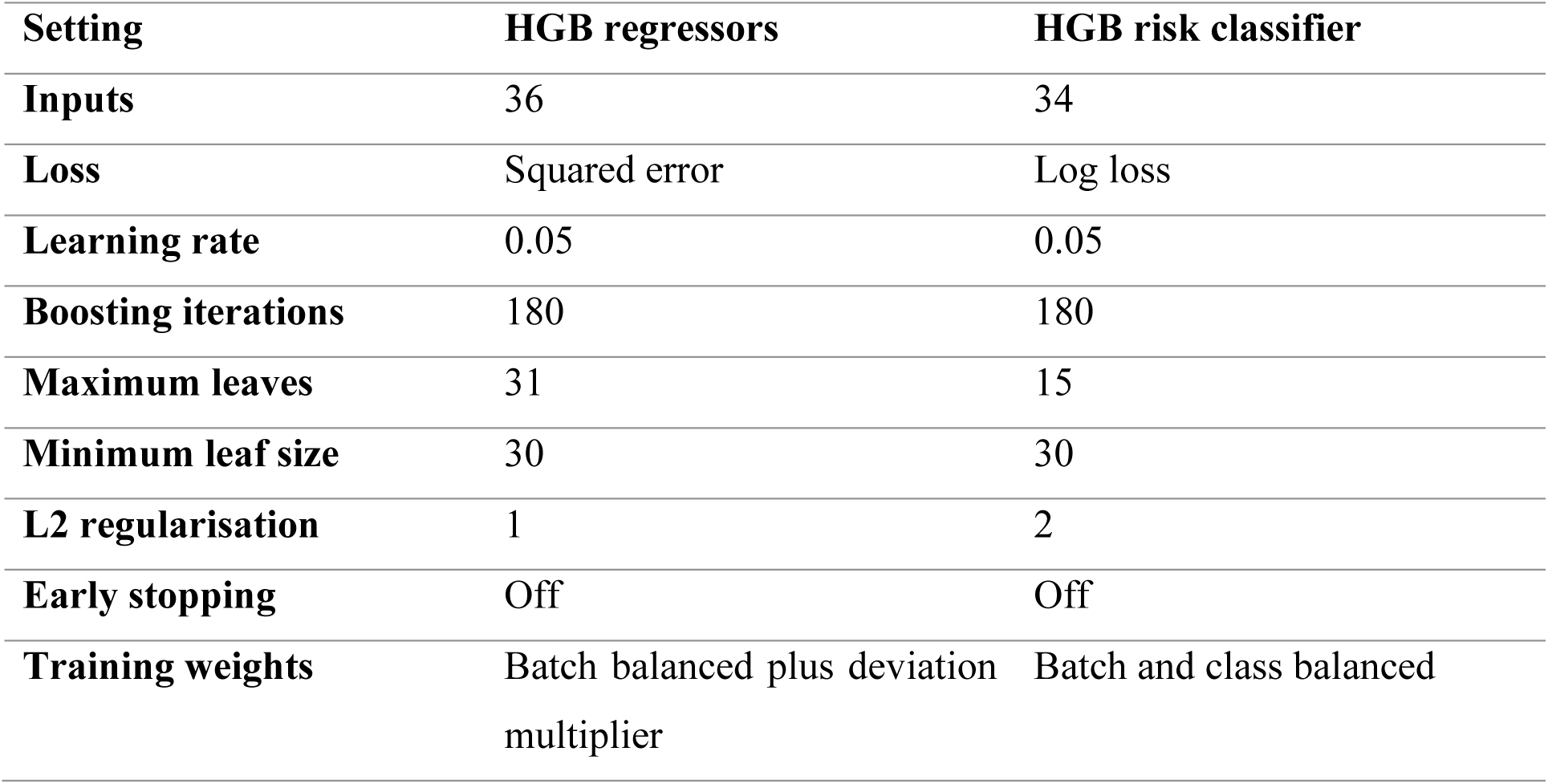
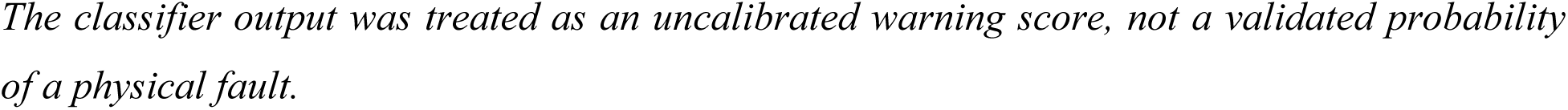
Settings used in the principal HGB components.

| Setting | HGB regressors | HGB risk classifier |
| --- | --- | --- |
| **Inputs** | 36 | 34 |
| **Loss** | Squared error | Log loss |
| **Learning rate** | 0.05 | 0.05 |
| **Boosting iterations** | 180 | 180 |
| **Maximum leaves** | 31 | 15 |
| **Minimum leaf size** | 30 | 30 |
| **L2 regularisation** | 1 | 2 |
| **Early stopping** | Off | Off |
| **Training weights** | Batch balanced plus deviation multiplier | Batch and class balanced |

The classifier output was treated as an uncalibrated warning score, not a validated probability of a physical fault.

### 3.7 Outer evaluation and statistical summaries

Normal-operation performance used five regime-balanced outer folds, each holding out 18 normal batches. The normal-only model was trained on the other 72 normal batches. The fault-inclusive model was trained on those same 72 normal batches plus all ten documented deviation batches; no normal test batch entered either fit. Deviation performance used ten leave-one-batch-out tests. The normal-only model was fitted to all 90 normal batches. For each held-out deviation batch, the fault-inclusive model was fitted to all 90 normal batches and the other nine deviation batches. Thus, every concentration prediction entering the main comparison came from a model that did not fit the tested batch. Holding out a batch does not guarantee that its physical deviation mechanism is absent from other training batches.

**Table 4:**
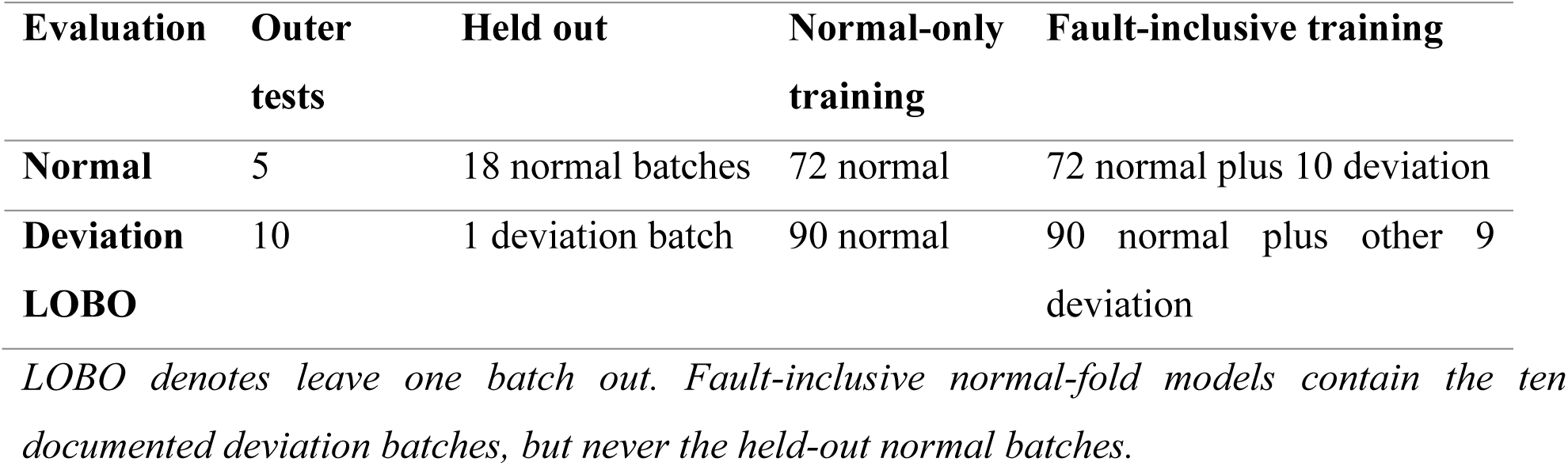
Outer complete-batch validation design.

MAE, RMSE and R² were calculated by pooling all held-out rows for each condition and separately for every batch. Pooled row metrics give longer batches more influence; per-batch metrics treat trajectories as the reporting units. The difference for each batch was fault-inclusive RMSE minus normal-only RMSE, so a negative value indicates improvement. Ten thousand bootstrap samples of paired complete-batch differences were drawn with replacement using seed 42 (Efron, 1979). Models were not refitted within the bootstrap. The resulting percentile intervals summarize the observed batch collection and are not formal generalization guarantees.

### 3.8 Reliability analysis

A separate HGB classifier used 34 features, omitting elapsed time and cumulative feed. Positive labels began at the first recorded fault activity and included all later rows in the documented deviation batches; normal rows and pre-onset rows were negative. Training sample weights balanced total contributions from batches and from positive and negative classes. A score of at least 0.5 generated a warning. The output was treated as an uncalibrated warning score, not a probability of a physical fault or a root-cause diagnosis. Performance was measured only from outer-held-out classifier predictions.

The final OOD detector was a separate descriptive deployment diagnostic. Up to 250 evenly spaced rows from each of the 90 normal batches produced 22,500 reference rows. Median imputation, standardization, PCA retaining at least 95% variance and a 400-tree Isolation Forest were fitted to that reference set. The negative of score_samples was used so larger values indicated greater unusualness, and the fitted-reference 99th percentile defined the warning threshold. This detector was not cross-fitted; its final scores therefore do not provide independent OOD accuracy on the development data.

Empirical concentration ranges blended the normal- and deviation-specific 90th-percentile absolute-error radii according to the fault-risk score. Ranges were widened by a factor of 1.25 for OOD rows and truncated at zero on the lower side. Because the radii were estimated from held-out residuals but the blending rule was not subjected to a separate coverage study, they are descriptive error ranges rather than calibrated confidence, prediction or safety intervals.

### 3.9 Software and reproducibility

The main archived run used Python 3.13.15, NumPy 2.3.5, pandas 2.2.3 and scikit-learn 1.8.0. The earlier experiments used scikit-learn 1.6.1. Automated repository tests verify data integrity, selected calculations and file consistency but do not retrain the models. A separate numerical reproduction executed the scientific cells sequentially after a Jupyter-kernel startup limitation and compared regenerated outputs with the archive. It passed 239 checks, including all five normal and ten deviation outer tests, with maximum absolute numeric discrepancy 2 × 10⁻¹⁵. This confirms numerical agreement in that execution context, not portability to every environment or independent fermentation validation.

## 4 Results

### 4.1 Initial model selection and normal trained baseline

Random Forest achieved validation MAE 1.228 g/L, RMSE 2.034 g/L and R² 0.9600, outperforming Linear Regression and the median dummy (Table 5). After refitting on the 75 normal development batches, its pooled normal-test MAE, RMSE and R² were 1.331 g/L, 2.181 g/L and 0.9464. On the deviation stress test, the corresponding values worsened to 2.712 g/L, 4.321 g/L and 0.7403. Figure S2 shows that the deterioration included systematic deviations from the equality line rather than only a small number of isolated points.

**Table 5:** Initial model selection on the normal validation batches.

| Model | MAE g/L | RMSE g/L | R<sup>2</sup> |
| --- | --- | --- | --- |
| Dummy Median | 8.851 | 10.175 | -0.0000 |
| Linear Regression | 1.817 | 2.539 | 0.9378 |
| Random Forest | 1.228 | 2.034 | 0.9600 |

*Random Forest was selected by the lowest validation RMSE. These are pooled row metrics on the original 15-batch normal validation subset.*

Impurity-based Random Forest importance was concentrated in cumulative sugar feed and elapsed time, with values 0.5193 and 0.4013; together they accounted for 92.06% of total importance (Figure S3). Off-gas oxygen was the next feature at 0.0382. These importances reflect how this fitted forest split its available predictors. Correlation, encoded batch progress and model bias prevent their interpretation as causal effects or as a complete biological ranking.

### 4.2 Findings from the five diagnostic experiments

Across 25 repeated regime-stratified complete-batch Random Forest fits, mean pooled RMSE was 2.050 g/L with standard deviation 0.286 and range 1.543–2.787 g/L. Mean MAE was 1.232 g/L and mean R² was 0.9568. No fold produced a negative median batch R². Figure S4 and Table S3 show the full fold distribution. Repetition reduced dependence on one partition but did not create new biological batches.

Removing time alone increased normal RMSE by 0.54% and reduced deviation RMSE by 1.88%, whereas removing cumulative feed increased normal and deviation RMSE by 1.59% and 0.99%. Removing both increased normal RMSE by 11.53% and deviation RMSE by 2.44% (Table 6 and Figure S5). The weakness under deviations therefore did not disappear when these two explicit progress variables were removed.

**Table 6:** Random Forest ablation of time and cumulative feed.

| Input set | Input<br>s | Normal RMSE | Deviation<br>RMSE | Deviation<br>change |
| --- | --- | --- | --- | --- |
| **All inputs** | 36 | 2.179 | 4.322 | +0.00% |
| **Without time** | 35 | 2.190 | 4.241 | -1.88% |
| **Without cumulative feed** | 35 | 2.213 | 4.365 | +0.99% |
| **Without both** | 34 | 2.430 | 4.427 | +2.44% |

*RMSE is in g/L. Change is relative to the all-input fit in this ablation experiment. The experiment does not remove every possible batch-progress proxy.*

The earlier Random Forest phase analysis showed pooled MAE of 0.112 g/L before the first recorded activity, 2.170 g/L within the first-to-last activity window and 4.708 g/L after the last activity. Mean per-batch MAE was 0.061, 1.485 and 4.721 g/L, respectively; the final phase included nine rather than ten batches. Figure S6 and Table S11 retain the batch-level values. Error growth coincided with later phases, but phase, time, target range and deviation activity were confounded.

HGB produced the lowest normal and deviation RMSE among the learned normal-trained comparators, at 2.030 and 4.190 g/L, but its deviation MAE of 2.768 g/L was slightly higher than the Random Forest value of 2.712 g/L. All learned model families showed a marked RMSE increase from normal to deviation batches (Table 7 and Figure S7). This result motivated a training-strategy comparison within HGB rather than the claim that one algorithm alone solved abnormal-operation prediction.

**Table 7:** Earlier normal trained model family comparison.

| **Model** | **Normal RMSE** | **Deviation RMSE** | **Normal R<sup>2</sup>** | **Deviation R<sup>2</sup>** |
| --- | --- | --- | --- | --- |
| **Dummy Median** | 9.647 | 9.916 | -0.0490 | -0.3678 |
| **Linear Regression** | 3.355 | 5.448 | 0.8731 | 0.5871 |
| **Random Forest** | 2.181 | 4.321 | 0.9464 | 0.7403 |
| **HistGradientBoosting** | 2.030 | 4.190 | 0.9535 | 0.7558 |

*RMSE is in g/L. The HGB comparator used 300 iterations and differs from the final 180-iteration HGB configuration.*

Early prefix-summary OOD detection flagged 2, 6, 6 and 4 of the ten deviation batches at 12, 24, 48 and 72 h. The corresponding normal-test counts were 0, 2, 1 and 0 of 15 (Table S4 and Figure S8). At 24 h, the association between OOD score and whole-batch Random Forest RMSE was heterogeneous (Figure S9); several high-error trajectories were not uniquely separated. The earlier 90% empirical ranges covered 84.89% of normal-test rows and 70.41% of deviation-test rows, with mean full widths 5.034 and 5.200 g/L (Table S5). These findings supported retaining warnings as auxiliary diagnostics rather than presenting them as dependable guarantees.

### 4.3 Principal normal-only and fault-inclusive comparison

Normal-operation performance was essentially unchanged between the two training strategies. Pooled normal RMSE was 1.9806 g/L for normal-only HGB and 1.9830 g/L for fault-inclusive HGB; MAE changed from 1.2095 to 1.2063 g/L and R² from 0.9607 to 0.9606. On held-out deviation batches, fault-inclusive training reduced pooled RMSE from 3.1946 to 2.5644 g/L, a 19.73% reduction, and MAE from 2.0763 to 1.4405 g/L, a 30.62% reduction. Fault R² increased from 0.8580 to 0.9085 (Table 8 and Figure 2). The pooled results weight rows, so longer batches contribute more.

**Figure 1:**
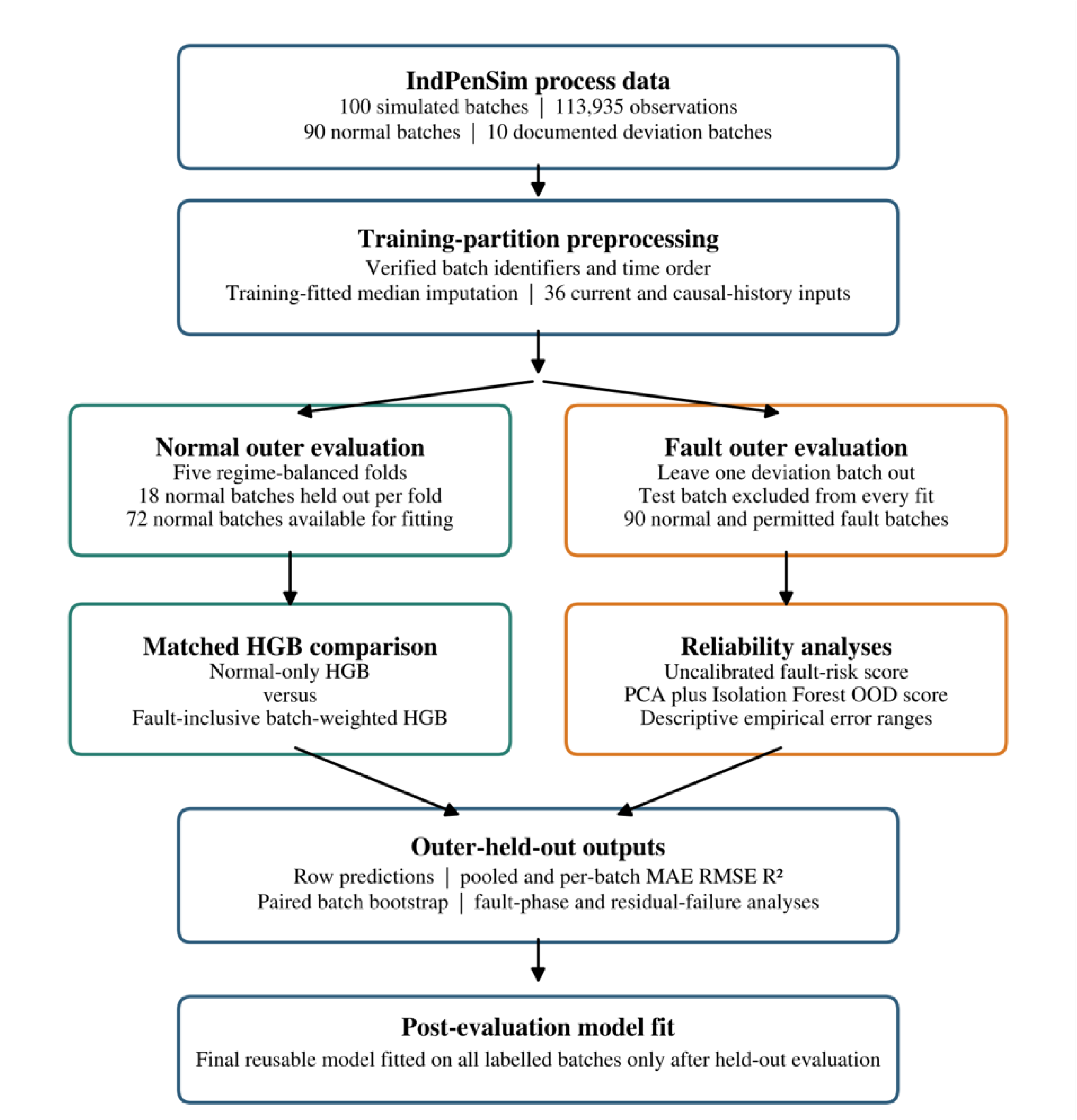
Complete batch evaluation workflow. Workflow of the principal analysis. Both training strategies use the same current-concentration target, 36 input definitions and HGB regressor settings. Normal evaluation holds out 18 complete normal batches per fold; deviation evaluation holds out one of the batches 91–100 at a time. Fault-inclusive fitting receives only the permitted batches. Imputation is fitted on each training partition. Paired errors use stored outer-held-out predictions, and the final reusable model is fitted only after evaluation. Holding out one batch does not establish the absence of a deviation mechanism in all training batches.

**Figure 2:**
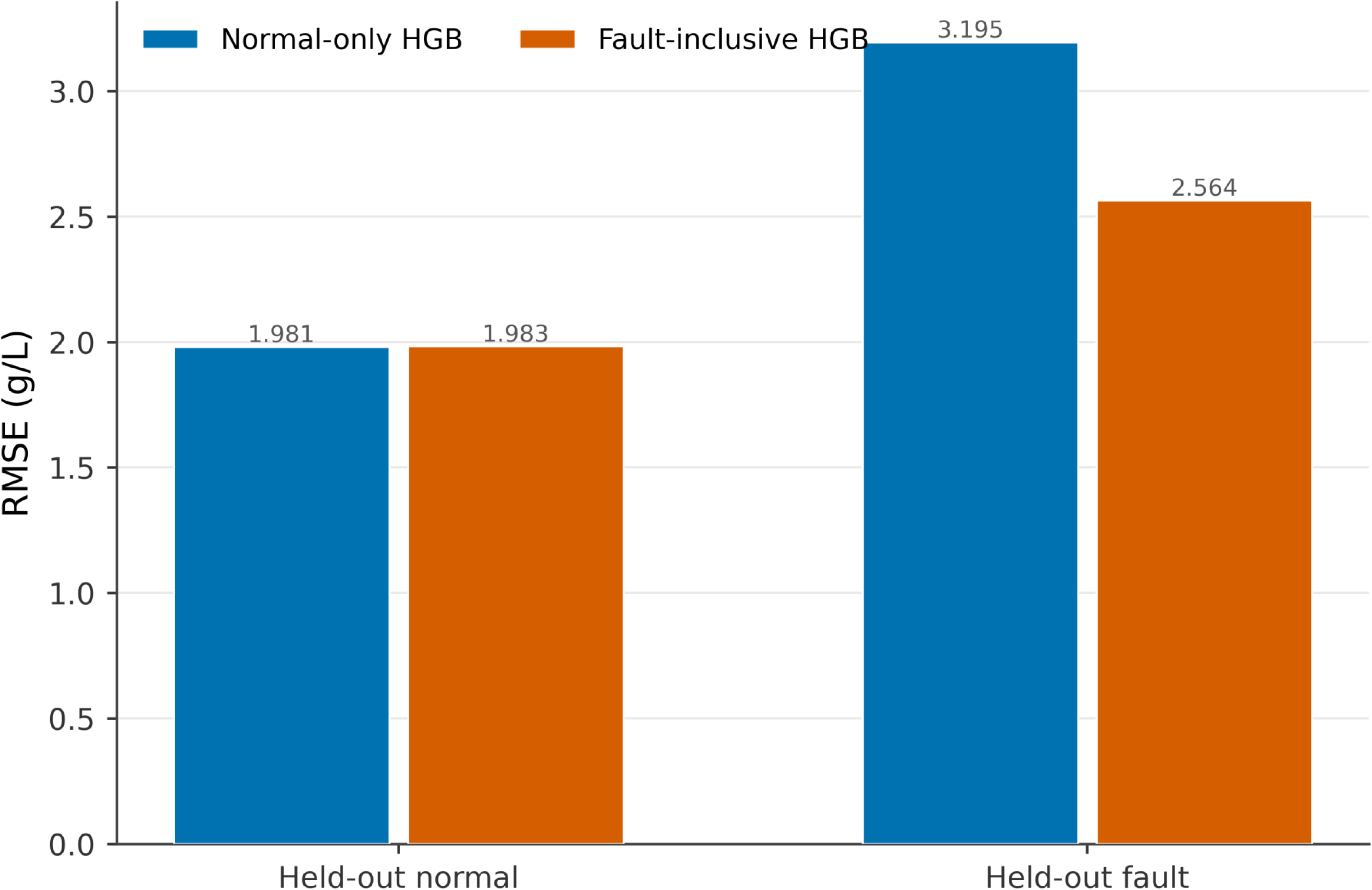
Pooled normal and deviation RMSE (Root Mean Square Error). Pooled RMSE for normal-only and fault-inclusive HGB under complete-batch evaluation. Normal results combine 102,410 rows from 90 held-out normal trajectories; deviation results combine 11,525 rows from ten leave-one-deviation-batch-out tests. Bars are point estimates, not means with confidence intervals. Deviation RMSE decreases from 3.1946 to 2.5644 g/L, while normal RMSE changes from 1.9806 to 1.9830 g/L. Longer batches exert more influence on these pooled metrics.

**Table 8:** Principal outer held out performance.

| Condition | Training strategy | MAE g/L | RMSE g/L | R<sup>2</sup> |
| --- | --- | --- | --- | --- |
| Normal | Normal-only HGB | 1.2095 | 1.9806 | 0.9607 |
| Normal | Fault-inclusive HGB | 1.2063 | 1.9830 | 0.9606 |
| Deviation | Normal-only HGB | 2.0763 | 3.1946 | 0.8580 |
| Deviation | Fault-inclusive HGB | 1.4405 | 2.5644 | 0.9085 |

*Metrics pool all outer-held-out rows within a condition. Display terminology is standardized as fault-inclusive HGB while archived identifiers remain unchanged for reproducibility.*

### 4.4 Batch level consistency

**Table 9:**
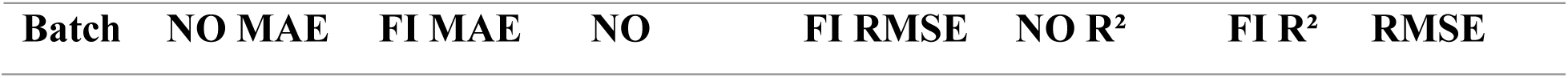
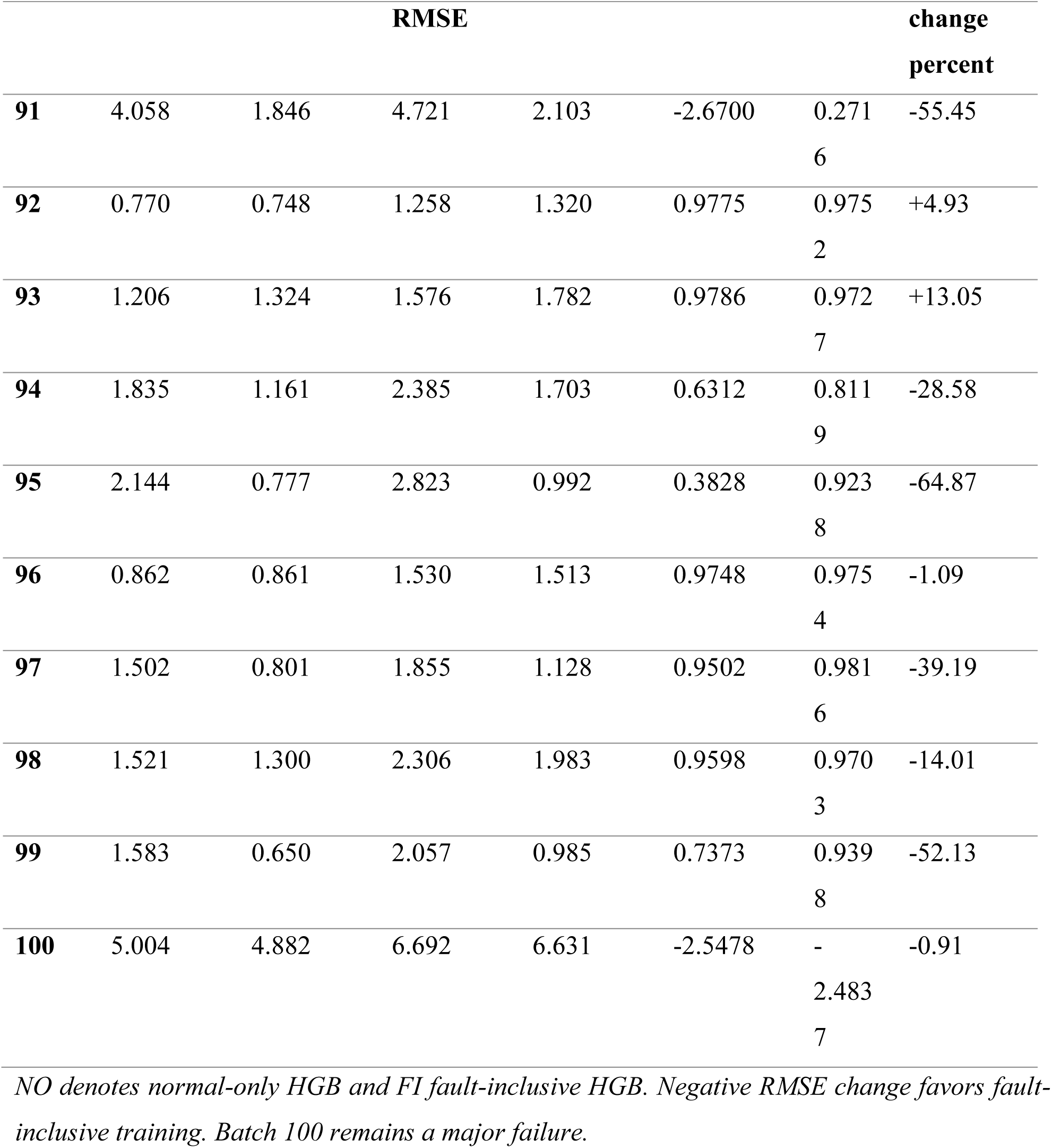
Held out performance for each documented deviation batch.

Fault-inclusive HGB reduced RMSE in eight of ten deviation batches (Figure 3). The largest proportional reductions occurred for batches 95, 91 and 99. Batches 92 and 93 worsened by 4.93% and 13.05%, respectively. Batch 100 changed only from 6.692 to 6.631 g/L and remained worse than a batch-mean predictor, as shown by R² −2.4837. Thus, the pooled improvement did not represent uniform reliability.

**Table 10:**
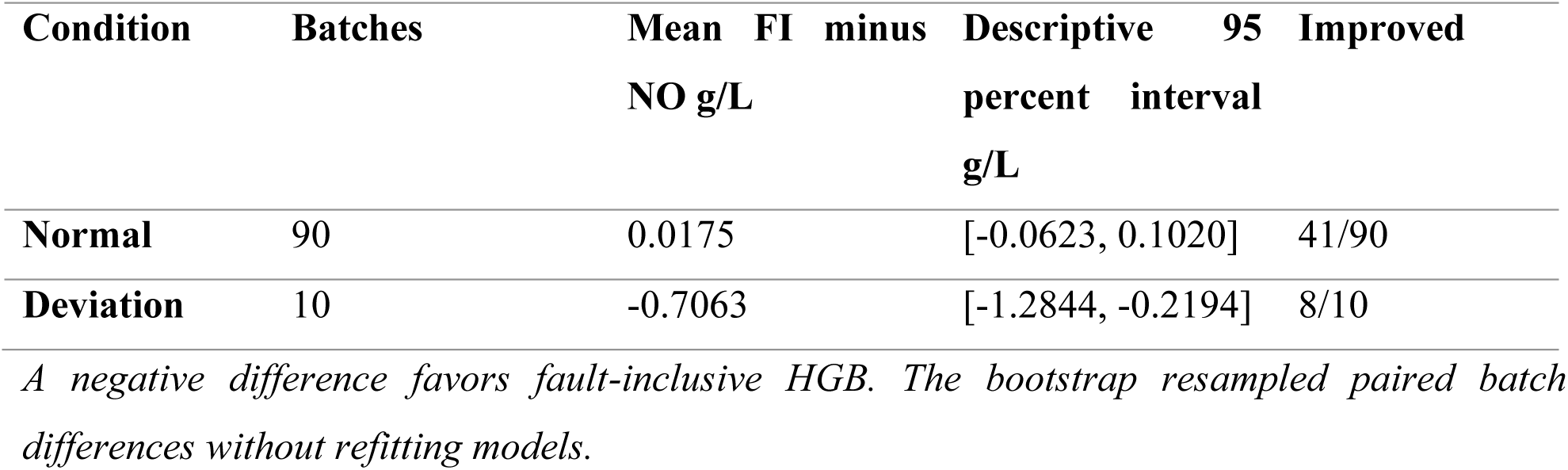
Paired complete-batch RMSE differences.

**Figure 3:**
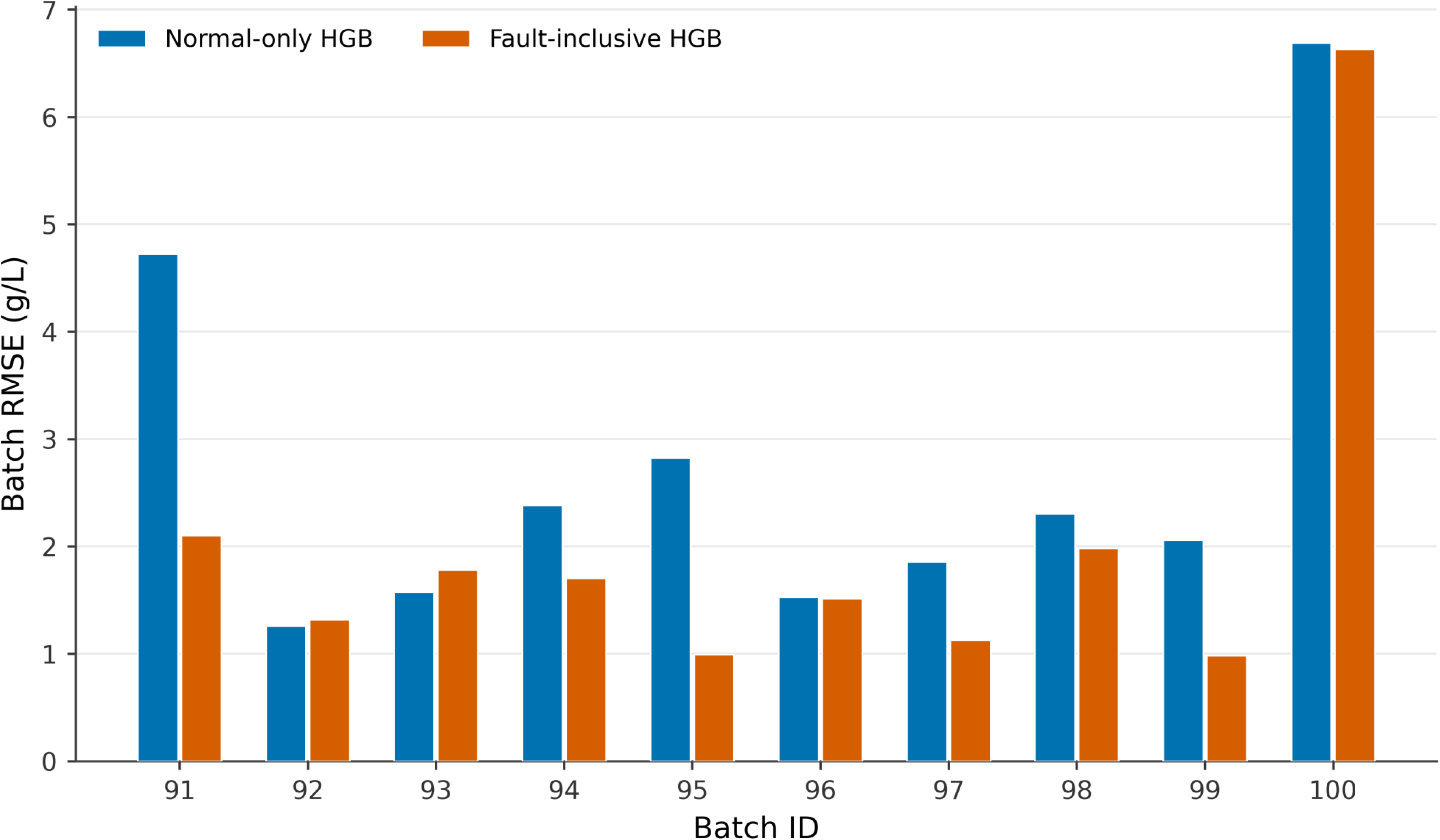
RMSE (Root Mean Square Error) in each held-out deviation batch. Per-batch RMSE for batches 91–100. The normal-only model is fitted to 90 normal batches. For each fault-inclusive bar, the model is fitted to those normal batches and the other nine deviation batches; the tested batch is excluded. Eight batches improve, batches 92 and 93 worsen, and batch 100 remains poorly estimated. Each bar represents a single held-out prediction trajectory rather than a replicated experimental mean.

**Figure 4:**
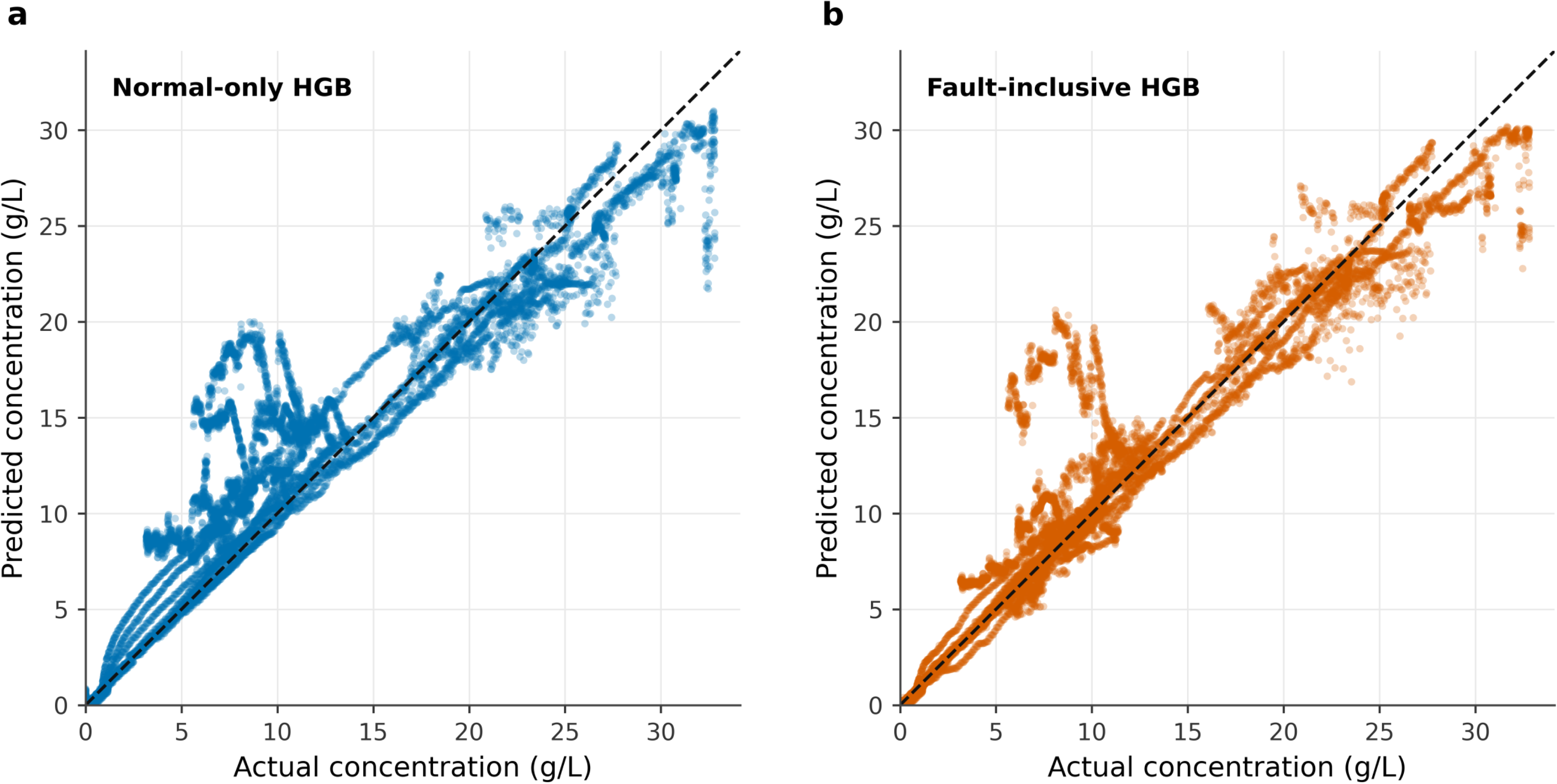
Actual and predicted deviation batch concentrations. Actual-versus-predicted penicillin concentration for all 11,525 outer-held-out deviation-batch observations. The dashed diagonal denotes equality. Points within each batch are serially dependent and are not independent experiments. Fault-inclusive fitting improves agreement for many observations, but substantial overprediction at low actual concentrations remains. The plot evaluates current concentration, not a future forecast horizon.

The mean paired fault-batch difference was −0.7063 g/L, with a descriptive bootstrap interval from −1.2844 to −0.2194 g/L. For normal batches, the mean difference was +0.0175 g/L and the interval crossed zero, from −0.0623 to +0.1020 g/L. Forty-one of 90 normal batches and eight of ten deviation batches improved. The fault interval is encouraging for this benchmark, but only ten potentially related deviation trajectories were available, and the leave-one-batch-out models shared most training data

### 4.5 Phase specific deviation performance

**Table 11:**
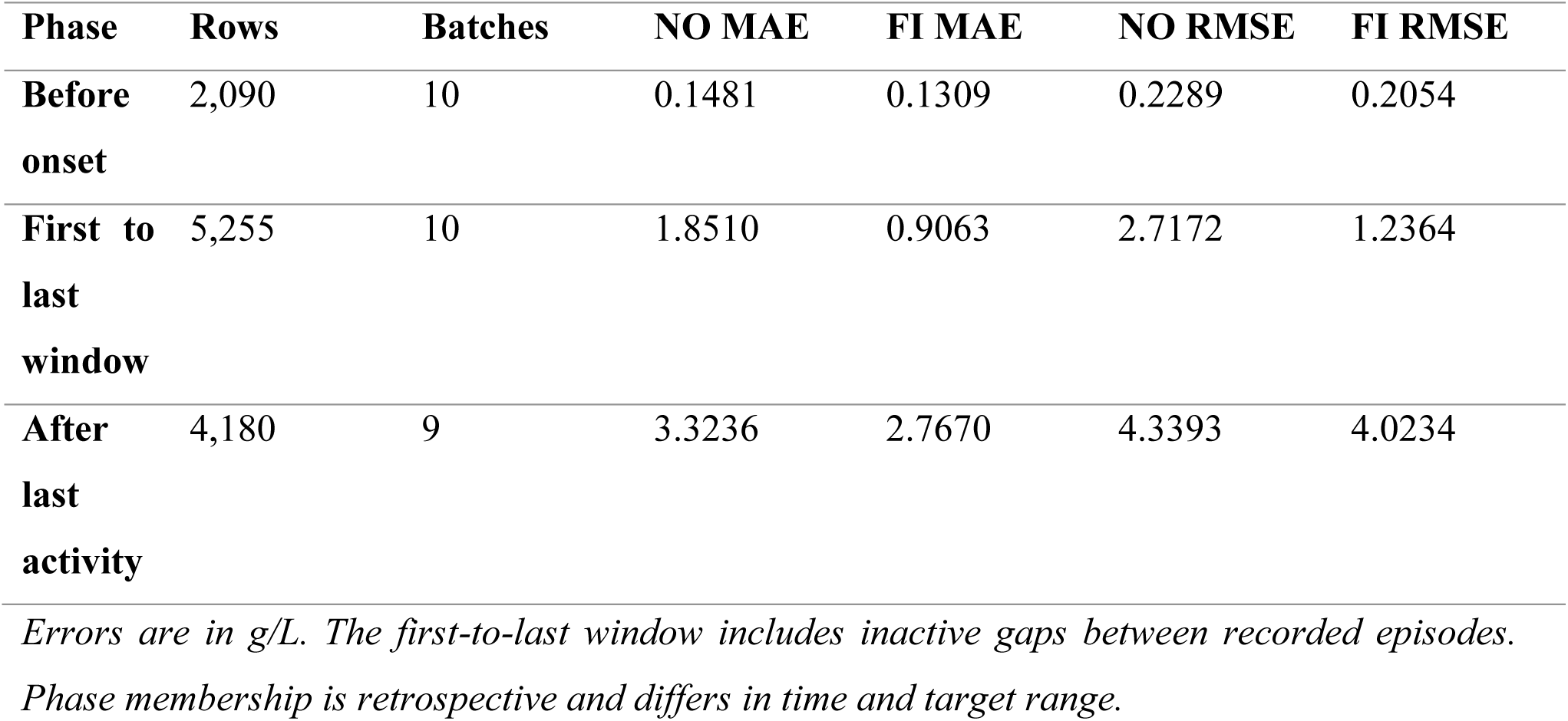
Principal HGB performance by retrospective fault reference phase.

Before the first recorded fault activity, both models were highly accurate and fault-inclusive RMSE was 0.2054 versus 0.2289 g/L. Within the first-to-last reference window, RMSE decreased from 2.7172 to 1.2364 g/L. After the last activity, the reduction was smaller, from 4.3393 to 4.0234 g/L. The largest improvement therefore occurred during the recorded window, whereas the difficult late-stage trajectories remained only partly corrected.

### 4.6 Warning scores OOD diagnostics and empirical ranges

**Table 12:** Outer held out fault risk warnings at score 0.5.

| Evaluation group | Rows | Warned rows | Warning rate |
| --- | --- | --- | --- |
| Normal rows | 102,41<br>0 | 14,189 | 13.86% |
| Pre-onset rows in deviation batches | 2,090 | 97 | 4.64% |
| Onset and after rows in deviation batches | 9,435 | 4,648 | 49.26% |

*The score is uncalibrated. The onset-and-after label includes rows after the last active reference. The within-fault-batch ROC AUC was 0.8434.*

The fault-risk classifier warned on 13.86% of held-out normal rows, 4.64% of pre-onset rows within deviation batches and 49.26% of onset-and-after rows. Its within-deviation-batch ROC AUC was 0.8434. Median scores were 0.0317 for normal rows, 0.0133 before onset and 0.4925 for onset and after rows (Figure 6). The overlap between groups explains why approximately half of affected rows were missed at the 0.5 threshold. Any-warning aggregation would have flagged all ten deviation batches but also 82 of 90 normal batches, which is unsuitable as a selective batch alarm.

**Figure 5:**
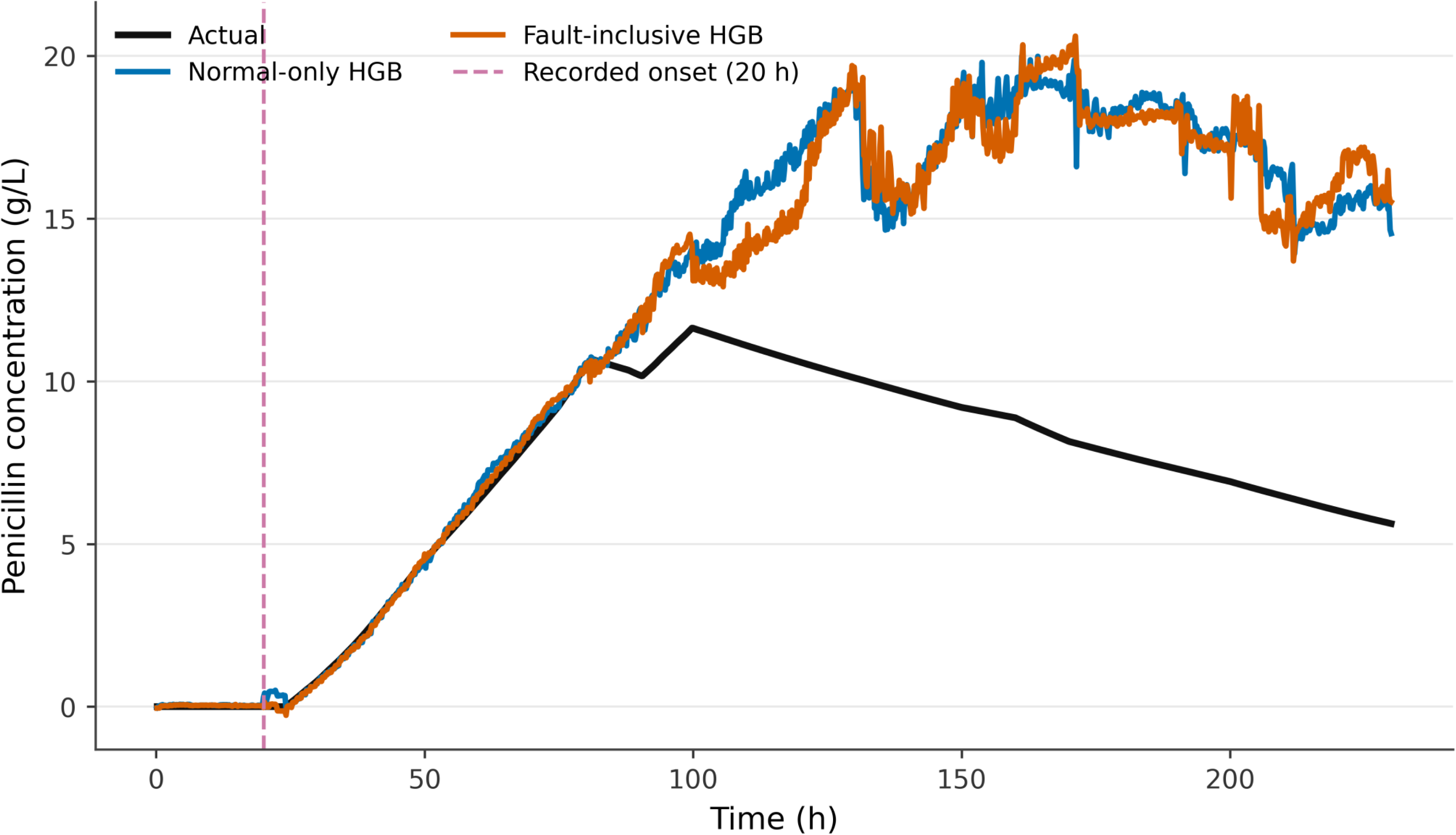
Batch 100 has an unresolved failure. Held-out concentration trajectory for batch 100. The black line is the simulated target; blue and orange lines are normal-only and fault-inclusive estimates. The vertical dashed line marks the first non-zero recorded fault reference at 20 h, not a model-detected onset. Neither regressor was fitted on batch 100. Both models increasingly overestimate the later low-producing trajectory. Fault-inclusive RMSE is 6.631 g/L, and R² is −2.4837; the last recorded fault activity occurs at 110 h.

**Figure 6:**
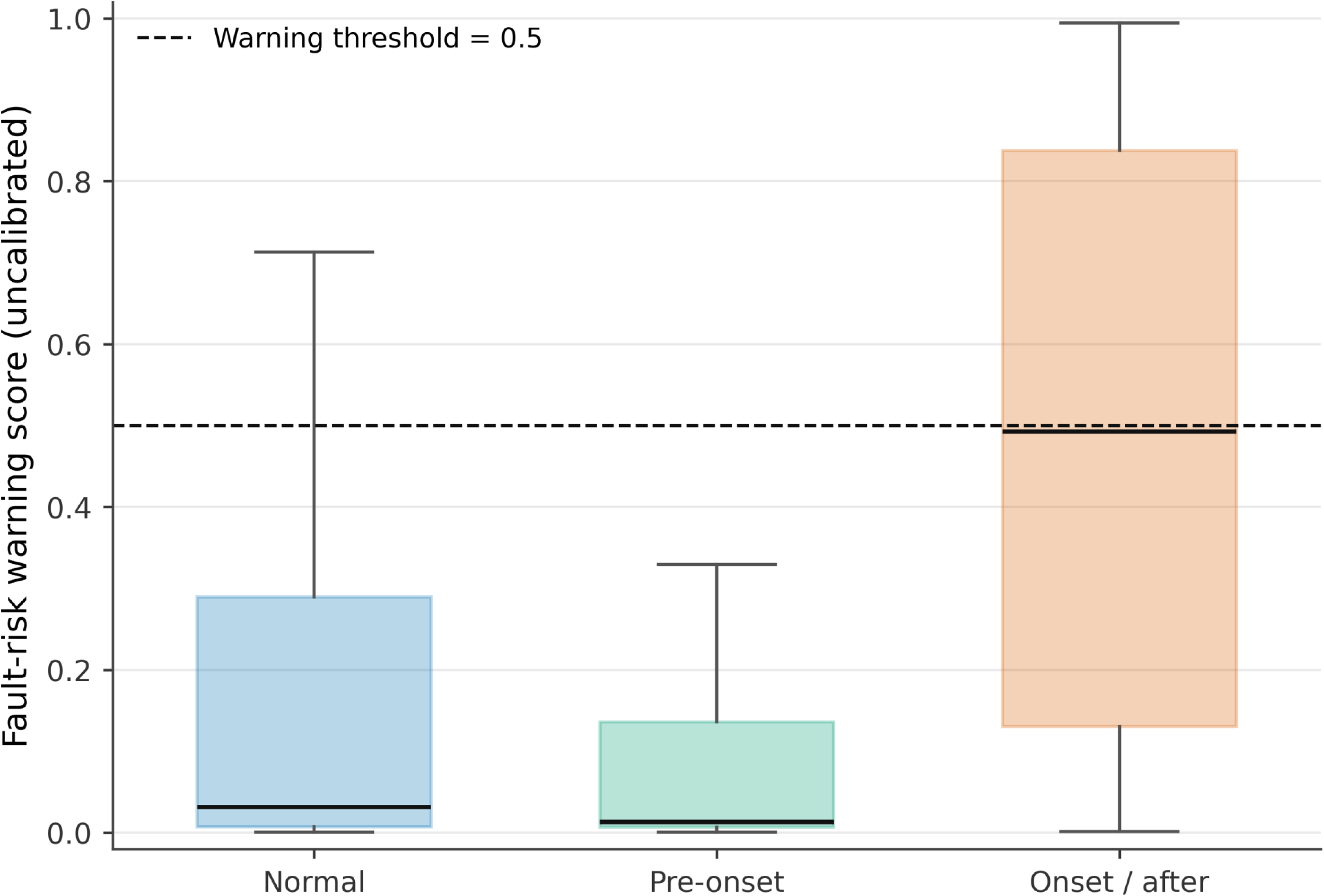
Outer-held-out fault-risk score distributions. Distributions of fault-risk scores in normal rows, pre-onset deviation-batch rows and onset-and-after deviation-batch rows. Boxes span the 25th–75th percentiles, centre lines show medians, and whiskers show the 5th–95th percentiles; points outside those whiskers are omitted. The dashed line is the 0.5 warning threshold. Scores are outer-held-out classifier outputs, are not calibrated physical-fault probabilities and do not correct the concentration estimate.

The final OOD reference used 15 principal components to retain 95.005% of variance, and its 99th-percentile score threshold was 0.626829. Fault-batch row warning rates were low to moderate and batch-specific (Table S9); batch 100 had an OOD rate of only 4.61%, despite being the largest concentration-prediction failure. OOD therefore did not provide a reliable surrogate for error. The normal and deviation empirical absolute-error radii were 3.0398 and 3.1164 g/L. Because the final blending and OOD-widening rule was not independently calibrated, these values were retained as descriptive deployment diagnostics only.

### 4.7 Residual failure in batch 100

Batch 100 illustrates the limitation hidden by pooled metrics. Its mean actual concentration was 6.931 g/L, whereas the mean fault-inclusive estimate was 11.790 g/L. At 230 h, the actual concentration was 5.626 g/L and the fault-inclusive estimate 15.511 g/L. Both models increasingly overpredicted the later low-producing trajectory (Figure 5). The modest RMSE change of −0.91% therefore did not represent a practically resolved failure, and neither the risk classifier nor OOD detector supplied a sufficiently dependable safeguard.

## 5 Discussion

The central result is a matched within-family comparison: fault-inclusive, batch-weighted HGB reduced pooled held-out deviation RMSE by 19.73% and improved eight of ten documented deviation batches, while pooled normal RMSE changed by only 0.12%. This supports the practical intuition that a regressor trained only on routine trajectories may benefit from representative abnormal examples. The result is stronger than a random row split because every reported prediction came from a model that did not fit the tested batch.

The study also shows why pooled accuracy is insufficient. Two deviation batches worsened, and batch 100 remained severely misestimated. Negative batch-level R² means that the trajectory was predicted worse than simply using its own mean target, although that mean would not be available prospectively. The adverse cases are not exceptions to be removed; they define the boundary of the current model.

The findings are consistent with the broader literature that treats maintenance and changing operating conditions as central soft-sensor problems (Brunner et al., 2021; Kadlec et al., 2009; Kadlec et al., 2011; Luttmann et al., 2012). They also extend the IndPenSim evidence that process deviations can degrade a normal-trained soft sensor and may require monitoring or retraining (Acosta-Pavas et al., 2024; Metcalfe et al., 2025). The specific contribution is not a novel boosting algorithm. It is the paired complete-batch evaluation of a fixed HGB family under two training strategies, together with transparent residual-failure analysis. Work outside IndPenSim helps clarify what the present method does not do. The adaptive ensemble of Siegl et al. reacts to sensor reliability (Siegl et al., 2022), and the online Gaussian-process ensemble of Jin et al. updates a model for time-varying batches (Jin et al., 2015). The present regressor is fitted offline and does not adapt during a held-out trajectory. Rivera et al. and Del Hierro et al. used physical fermentation data (Del Hierro et al., 2026; Rivera et al., 2024), including independent-batch validation in the latter study, whereas all observations here arise from a simulator. Liu et al. focused on sensor-failure tolerance (Liu et al., 2024), while this benchmark contains documented process deviations. These differences preclude direct error ranking and indicate distinct routes for future validation.

The five earlier experiments reduce several simple alternative explanations. Repeated complete-batch validation showed that normal Random Forest performance was not an artefact of one partition. Removing time and cumulative feed did not remove the deviation error. Normal-trained Linear Regression, Random Forest and HGB all deteriorated on the stress test, so the problem was not unique to one algorithm. Phase analysis showed that late trajectories were especially difficult, and early OOD or empirical ranges did not provide complete protection. These experiments motivated the final comparison but are not independent replications because they reuse the same benchmark and informed model development.

Fault-inclusive improvement was largest inside the recorded first-to-last activity window. This may indicate that the permitted deviation examples helped HGB represent some altered input– target relationships while reference activity was present. It does not prove that the model learned a physical fault mechanism. Phase is entangled with elapsed time, product concentration, batch duration and the particular simulated scenarios. Mechanism-held-out tests are needed to determine whether benefit transfers to a genuinely new abnormal condition. The contrast among batches suggests that diversity of abnormal training examples matters. Strong gains in batches 91, 95 and 99 coexisted with worsening in 92 and 93 and near-failure in 100. A future study should identify the documented deviation types, construct mechanism-level groups and hold out every batch belonging to one mechanism. A same-weight comparison would also separate fault inclusion from the selected multiplier of three.

The risk score ranked onset-and-after observations above pre-onset observations reasonably well in aggregate, but its threshold detected only 49.26% of affected rows and produced warnings on 13.86% of normal rows. The ROC AUC of 0.8434 is a ranking statistic inside the fault batches, not evidence that the score is calibrated or that it distinguishes all normal production from all faults. Probability calibration and precision-recall analysis would be necessary before interpreting threshold warnings operationally (Guo et al., 2017; Saito & Rehmsmeier, 2015).

OOD detection answered a different question: whether input combinations resembled the fitted normal reference. An unusual normal batch may be flagged, and a harmful process deviation may remain within the normal input distribution. Batch 100 demonstrates the latter possibility. More generally, uncertainty can degrade under distribution shift (Hüllermeier & Waegeman, 2021; Ovadia et al., 2019). The empirical ranges and OOD flag should therefore be treated as cautionary metadata, not a guarantee that the concentration estimate is safe to use. For deployment, the most defensible output is a bundle containing the concentration estimate, a clearly labelled uncalibrated risk score, an OOD score, an empirical range and provenance for the fitted model. A human operator would still need process context and laboratory confirmation. The current evidence does not justify autonomous control, release decisions or safety-critical use.

The results in this work can be interpreted in the context of digital-twin-enabled bioprocess monitoring. A complete bioprocess digital twin requires more than a bioprocess simulator, e.g. it requires reliable estimates of the evolving process state from measurements available during the fermentation operation. Measuring the product concentration is particularly relevant as it may not be measurable at the same way as conventional process variables. The soft sensor evaluated here represents such a component, translating available process information into an estimate of current penicillin concentration. The comparison of the normal-only with fault-inclusive training shows that the state-estimation layer had decreased reliability as the process diverged from nominal trajectories even when forecast performance under normal conditions was excellent. Fitting to a sample of representative deviation data resulted in a decreased pooled held-out deviation RMSE of 2.5644 g/L versus an improvement in the normal-operation RMSE to essentially unchanged. By maintaining performance in the normal domain at a very high level, deviating trajectories may train data-driven state estimation to be more robust in the operating space.

This improvement would not be considered evidence of a fault-tolerant or fully autonomous digital twin. Two deviation batches increased in magnitude, and batch 100 showed steady degradation after fault inclusive training. The fault-risk classifier, OOD detector and empirical bounds also failed to consistently offer reliable protection against all significant prediction errors. These observations also reveal an important digital-twin design criterion: a state estimate should ideally be accompanied by information describing whether the current operating condition is represented by the model and whether the estimate itself can be trusted. A more comprehensive bioprocess digital-twin approach could expand on the existing architecture by incorporating beyond current-state estimation, uncertainty quantification, online intervention decision support, and interaction by including dynamic prediction or forecasting, online model adaptation, mechanistic or hybrid process modeling, mechanism-aware fault detection, and two-direction interaction with an operational fermentation process. Validation against independent experimental fermentations would also be necessary before the system could be considered a validated digital twin for operational use.

## 6 Limitations

First, the study uses one simulated benchmark. The 100 trajectories do not represent the diversity, sensor artefacts, biological variability or interventions present in physical production. Results may depend on the simulator’s equations and the selected export. Raman spectra were excluded, and the online availability of every process derived variable was not demonstrated.

Second, the ten deviation batches are a small and possibly related collection. Leave-one-batch-out evaluation prevents direct batch leakage but does not guarantee mechanism novelty. The same benchmark informed baseline development, diagnostic experiments, and the final design. No separate external dataset was reserved after model-development decisions.

Third, fault inclusion and a threefold fault weight changed together, and the multiplier was not tuned within nested validation. The design therefore estimates the performance of that combined strategy, not the isolated effect of adding fault data. Feature-usability screening was fixed before the 25 repeated Random Forest fits rather than repeated within each fold. Random Forest impurity importance is also biased by correlated predictors and variable split opportunities.

Fourth, row-level metrics are serially dependent and pooled metrics weight longer trajectories more. Per-batch reporting mitigates but does not eliminate this limitation. The paired bootstrap resamples only ten deviation-batch differences, and the fitted models share training data, so its interval is descriptive. R² can be unstable in batches with limited target variation and should be read with MAE, RMSE and trajectories.

Fifth, the risk classifier was not probability-calibrated, the final OOD detector was not cross-fitted, and the empirical ranges were not assessed on an independent calibration set after the final blending rule. The warning components are not causal diagnoses. Negative concentration estimates occurred in 686 normal-only and 710 fault-inclusive held-out rows, with minima −0.3164 and −0.2640 g/L. Predictions were not clipped for metric calculation; physical non-negativity could be incorporated in future modelling.

Finally, numerical reproduction in a documented environment is not the same as independent scientific replication. The repository tests do not retrain models, and the sequential-cell reproduction was adapted to an infrastructure limitation. A clean end-to-end notebook run on a second system, a frozen release and physical-fermentation validation remain priorities.

## 7 Conclusions

Normal-trained penicillin soft sensors performed well on routine IndPenSim trajectories but showed substantial deterioration on documented process-deviation batches. Repeated complete-batch validation, progress-feature ablation, phase analysis, model-family comparison and early reliability experiments showed that this weakness was not explained by one split, two explicit progress variables or Random Forest alone.

Within the principal matched HGB comparison, fault-inclusive batch-weighted training reduced held-out deviation RMSE from 3.195 to 2.564 g/L and improved eight of ten batches, while normal RMSE remained approximately 1.98 g/L. Improvement was strongest during the recorded fault-reference window. Large late-stage errors persisted, especially in batch 100, and the warning components missed important failures.

The supported claim is therefore limited but useful: representative deviation data improved average current-concentration estimation within this simulated benchmark. Generalization to unseen deviation mechanisms, calibrated reliability and physical fermentation deployment has not been established. Mechanism-held-out testing, weight ablation, external experimental batches and prospectively defined reliability criteria are the most important next steps. From a digital-twin perspective, this proof of concept demonstrates the potential of fault-inclusive soft sensing as a proposed element of a true state-estimator for monitoring simulated bioprocesses outside nominal operating conditions. To achieve a full simulated bioprocess digital twin, component implementation must be advanced with integration into a forecasting, calibrated reliability assessment, online adaptation and inclusion of experimental fermentation data.

## Data availability

The public IndPenSim dataset is available from Mendeley Data, version 1, at https://doi.org/10.17632/pdnjz7zz5x.1 (Goldrick, 2019) under the stated Creative Commons Attribution 4.0 licence. This work used process-variable exports and did not use the 2,200 Raman columns. Split definitions, batch identifiers and checksums are recorded in the repository data manifest. The supplementary material documents the full analytical trail supporting the main comparison, including features, batch splits, validation folds, OOD and empirical-range analyses, fault windows, reproduction checks, fault-batch diagnostics, and the output inventory. Publication figures are paired with original code-output images containing the same evidence and should not be treated as independent analyses. Full row-level predictions remain in the repository rather than being reproduced in the supplement.

## Code availability

Analysis notebooks processed split definitions, held-out predictions, result tables and figures are maintained at https://github.com/abdulbasitbehlim/IndPenSim-penicillin-soft-sensor.

## Author contributions

Abdul Basit Behlim (ABB): conceptualization, methodology, investigation, software, validation, formal analysis, data curation, visualization and writing - original draft.

Atharva Tilewale (AT): project administration, writing - review and editing

Dhaval Patel (DP): supervision, project administration, resources, writing - review and editing.

## Acknowledgements

The authors further acknowledge the Gujarat Biotechnology University, Department of Science & Technology (DST), Government of Gujarat) for providing the necessary infrastructure facilities.

## Declaration of Generative AI and AI-assisted technologies in the writing process

During the preparation of this work the authors have used ChatGPT (GPT-5; https://chatgpt.com/) in order to improve the readability and language of the manuscript. After using this tool/service, authors have reviewed and edited the content as needed and take full responsibility for the content of the published article.

---

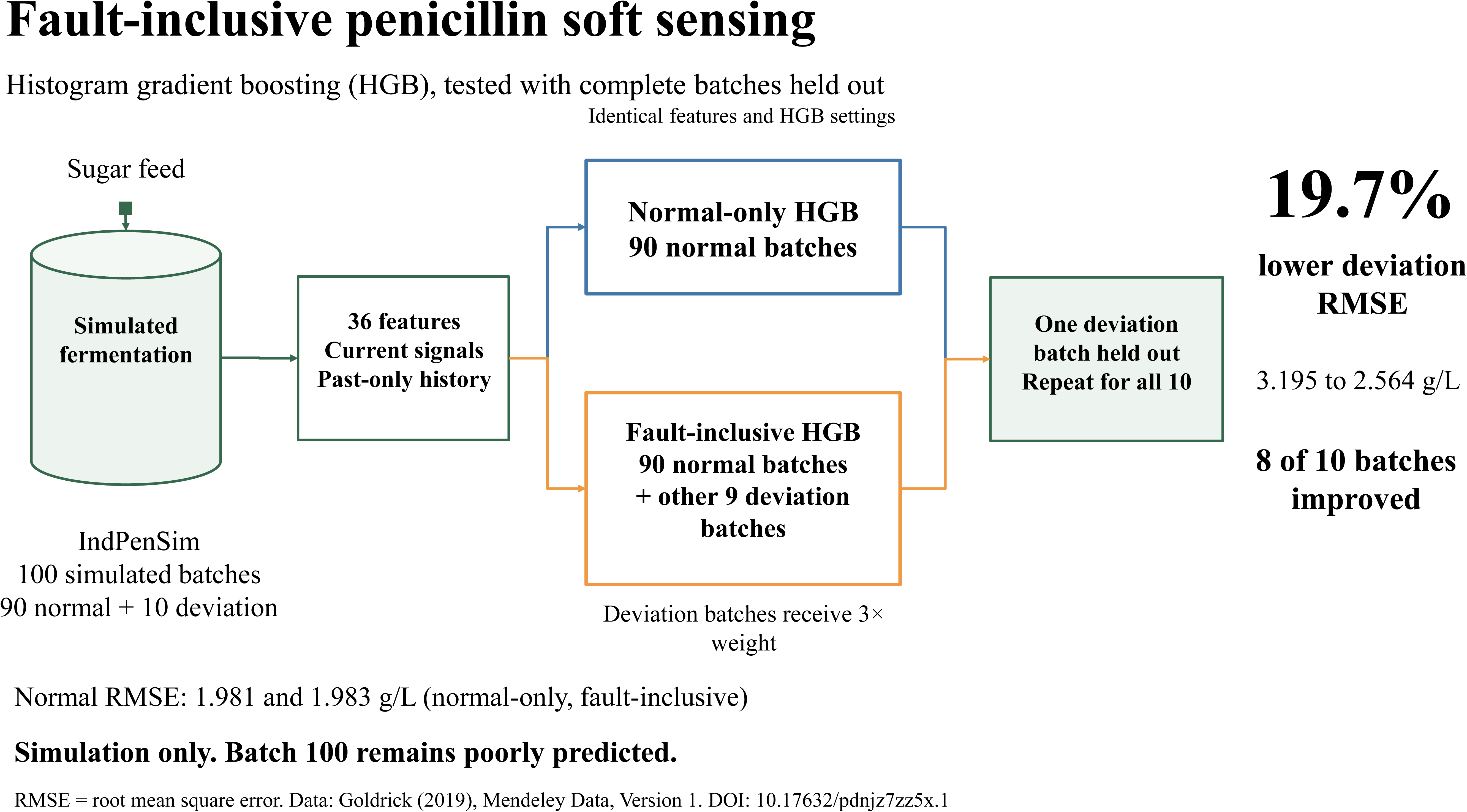

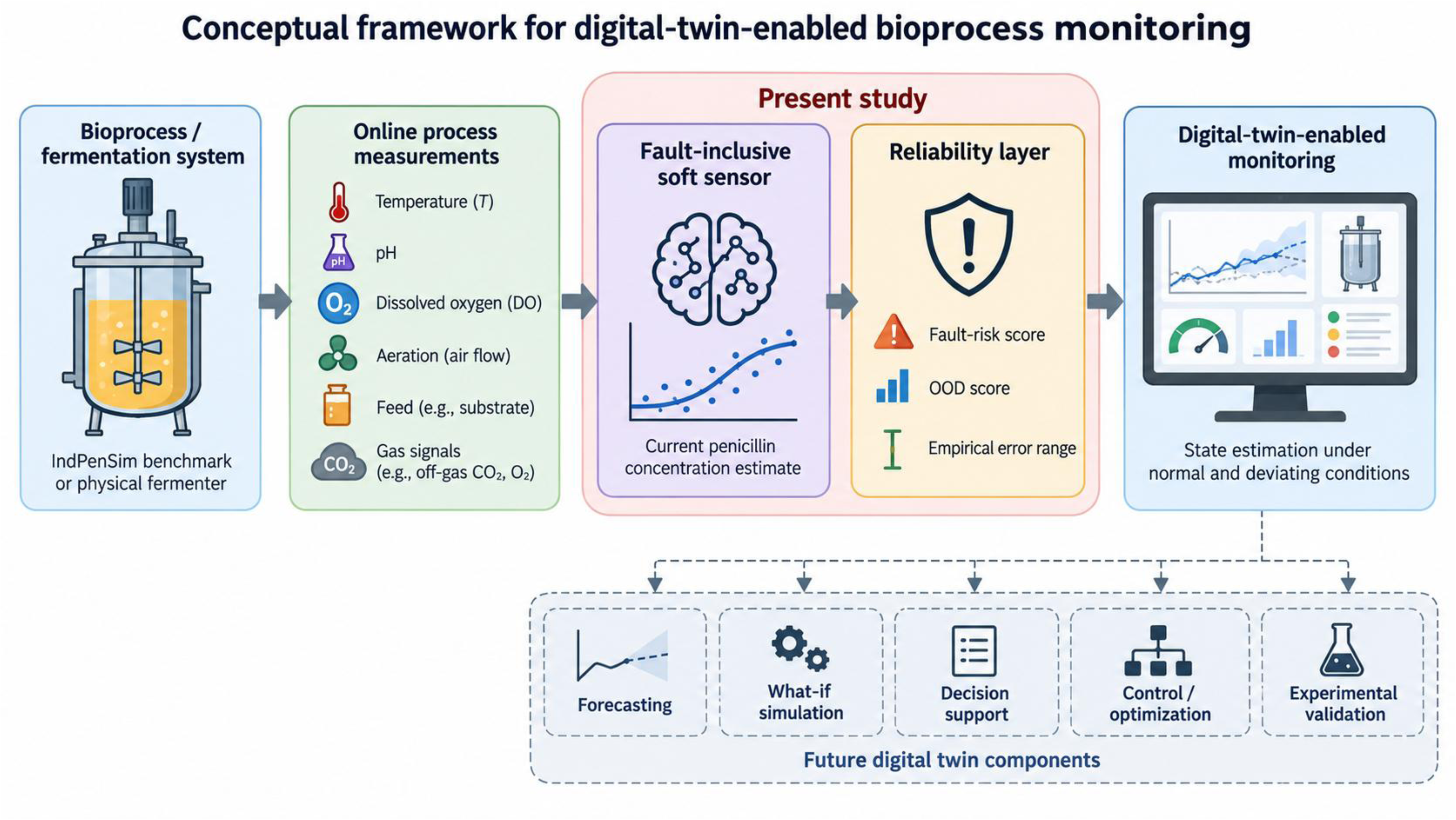

