## Supplemental Figures for "Towards Digital-Twin-Enabled Bioprocess Monitoring: Fault-Inclusive Soft Sensing of Penicillin Concentration Under Process Deviations"

**Supplementary information**

**Table S1 Exact regression feature names**

| Number | Saved feature name | Family | Risk and OOD input |
|---|---|---|---|
| 1 | Time (h) | Current or base | No |
| 2 | Aeration rate(Fg:L/h) | Current or base | Yes |
| 3 | Sugar feed rate(Fs:L/h) | Current or base | Yes |
| 4 | Acid flow rate(Fa:L/h) | Current or base | Yes |
| 5 | Base flow rate(Fb:L/h) | Current or base | Yes |
| 6 | Heating/cooling water flow rate(Fc:L/h) | Current or base | Yes |
| 7 | Heating water flow rate(Fh:L/h) | Current or base | Yes |
| 8 | Water for injection/dilution(Fw:L/h) | Current or base | Yes |
| 9 | Air head pressure(pressure:bar) | Current or base | Yes |
| 10 | Dissolved oxygen concentration(DO2:mg/L) | Current or base | Yes |
| 11 | Vessel Volume(V:L) | Current or base | Yes |
| 12 | Vessel Weight(Wt:Kg) | Current or base | Yes |
| 13 | pH(pH:pH) | Current or base | Yes |
| 14 | Temperature(T:K) | Current or base | Yes |
| 15 | Generated heat(Q:kJ) | Current or base | Yes |
| 16 | carbon dioxide percent in off-gas(CO2outgas:%) | Current or base | Yes |
| 17 | PAA flow(Fpaa:PAA flow (L/h)) | Current or base | Yes |
| 18 | Oil flow(Foil:L/hr) | Current or base | Yes |
| 19 | Oxygen Uptake Rate(OUR:(g min^{-1})) | Current or base | Yes |

| Number | Saved feature name | Family | Risk and OOD input |
|---|---|---|---|
| 20 | Oxygen in percent in off-gas(O2:O2 (%)) | Current or base | Yes |
| 21 | Dissolved oxygen concentration(DO2:mg/L)_lag1 | Within batch history | Yes |
| 22 | Dissolved oxygen concentration(DO2:mg/L)_difference1 | Within batch history | Yes |
| 23 | Dissolved oxygen concentration(DO2:mg/L)_previous5_mean | Within batch history | Yes |
| 24 | Sugar feed rate(Fs:L/h)_lag1 | Within batch history | Yes |
| 25 | Sugar feed rate(Fs:L/h)_difference1 | Within batch history | Yes |
| 26 | Sugar feed rate(Fs:L/h)_previous5_mean | Within batch history | Yes |
| 27 | Temperature(T:K)_lag1 | Within batch history | Yes |
| 28 | Temperature(T:K)_difference1 | Within batch history | Yes |
| 29 | Temperature(T:K)_previous5_mean | Within batch history | Yes |
| 30 | pH(pH:pH)_lag1 | Within batch history | Yes |
| 31 | pH(pH:pH)_difference1 | Within batch history | Yes |
| 32 | pH(pH:pH)_previous5_mean | Within batch history | Yes |
| 33 | Aeration rate(Fg:L/h)_lag1 | Within batch history | Yes |
| 34 | Aeration rate(Fg:L/h)_difference1 | Within batch history | Yes |
| 35 | Aeration rate(Fg:L/h)_previous5_mean | Within batch history | Yes |
| 36 | Cumulative_Sugar_Feed | Cumulative feed | No |

*Names preserve exported spelling and units. The 34 reliability inputs omit elapsed time and cumulative feed. Cumulative sugar feed has units of litres of feed because no concentration conversion was applied.*

**Table S2 Original baseline split membership**

| Subset | Data file | Rows | Batches | Batch IDs |
|---|---|---|---|---|
| Normal training | train_normal_60_batches.csv | 67820 | 60 | 1, 4, 6, 7, 8, 10, 11, 16, 17, 18, 19, 20, 21, 22, 23, 24, 25, 26, 27, 30, 31, 32, 33, 35, 36, 37, 38, 39, 41, 42, 44, 48, 49, 50, 51, 53, 54, 55, 57, 59, 61, 62, 63, 64, 65, 67, 72, 74, 75, 76, 79, 80, 81, 84, 85, 86, 87, 88, 89, 90 |
| Normal validation | validation_normal_15_batches.csv | 17510 | 15 | 3, 12, 13, 15, 29, 40, 43, 52, 56, 58, 68, 70, 73, 78, 82 |
| Normal test | test_normal_15_batches.csv | 17080 | 15 | 2, 5, 9, 14, 28, 34, 45, 46, 47, 60, 66, 69, 71, 77, 83 |
| Deviation stress test | test_fault_10_batches.csv | 11525 | 10 | 91, 92, 93, 94, 95, 96, 97, 98, 99, 100 |

*The three normal subsets contain equal numbers of recipe-driven, operator-controlled and advanced-process-control batches. Hashes are recorded in the repository manifest.*

**Table S3  All 25 repeated complete-batch Random Forest folds**

| Repeat | Fold | Test rows | MAE | RMSE | $R^2$ | Held out batch IDs |
|---|---|---|---|---|---|---|
| 1 | 1 | 20680 | 1.2808 | 2.1232 | 0.9578 | 8, 17, 23, 25, 26, 30, 32, 36, 39, 41, 42, 50, 63, 65, 72, 74, 75, 87 |
| 1 | 2 | 20245 | 1.3053 | 2.1269 | 0.9572 | 6, 11, 20, 21, 22, 27, 31, 33, 37, 48, 53, 55, 64, 76, 84, 85, 86, 89 |
| 1 | 3 | 19995 | 1.0246 | 1.6697 | 0.9714 | 1, 4, 7, 10, 16, 19, 35, 38, 49, 51, 57, 59, 61, 67, 79, 80, 81, 90 |
| 1 | 4 | 20685 | 1.2937 | 2.3103 | 0.9474 | 12, 13, 15, 18, 24, 29, 40, 44, 52, 54, 56, 58, 62, 68, 70, 78, 82, 88 |
| 1 | 5 | 20805 | 1.5154 | 2.4878 | 0.9283 | 2, 3, 5, 9, 14, 28, 34, 43, 45, 46, 47, 60, 66, 69, 71, 73, 77, 83 |
| 2 | 1 | 20395 | 1.0129 | 1.8278 | 0.9670 | 2, 6, 11, 14, 26, 28, 32, 34, 35, 38, 43, 56, 62, 63, 64, 66, 72, 90 |
| 2 | 2 | 20225 | 1.2287 | 1.9657 | 0.9591 | 7, 8, 13, 17, 19, 30, 36, 45, 46, 58, 59, 60, 61, 69, 71, 77, 83, 88 |
| 2 | 3 | 21635 | 1.4358 | 2.3365 | 0.9462 | 3, 9, 12, 24, 25, 29, 31, 33, 41, 44, 52, 53, 68, 78, 80, 82, 84, 86 |
| 2 | 4 | 20225 | 1.3937 | 2.2535 | 0.9481 | 10, 15, 18, 20, 22, 27, 37, 42, 48, 49, 51, 55, 65, 70, 73, 74, 79, 81 |
| 2 | 5 | 19930 | 0.9765 | 1.5987 | 0.9753 | 1, 4, 5, 16, 21, 23, 39, 40, 47, 50, 54, 57, 67, 75, 76, 85, 87, 89 |
| 3 | 1 | 20880 | 1.2165 | 2.1433 | 0.9477 | 1, 9, 10, 14, 16, 25, 33, 35, 39, 40, 42, 43, 61, 65, 73, 75, 79, 86 |
| 3 | 2 | 20200 | 0.9757 | 1.5430 | 0.9757 | 13, 17, 26, 27, 28, 30, 32, 37, 54, 55, 56, 58, 64, 78, 82, 88, 89, 90 |

| Repeat | Fold | Test rows | MAE | RMSE | $R^2$ | Held out batch IDs |
|---|---|---|---|---|---|---|
| 3 | 3 | 20260 | 1.2280 | 1.9985 | 0.9623 | 4, 5, 11, 12, 21, 23, 31, 34, 36, 50, 52, 59, 70, 71, 74, 76, 83, 85 |
| 3 | 4 | 21055 | 1.3613 | 2.2694 | 0.9510 | 3, 18, 19, 20, 22, 29, 38, 45, 48, 49, 51, 60, 63, 72, 77, 80, 81, 87 |
| 3 | 5 | 20015 | 1.2876 | 2.2422 | 0.9506 | 2, 6, 7, 8, 15, 24, 41, 44, 46, 47, 53, 57, 62, 66, 67, 68, 69, 84 |
| 4 | 1 | 20795 | 1.2802 | 2.0870 | 0.9558 | 3, 7, 14, 17, 28, 30, 32, 34, 41, 50, 52, 58, 61, 62, 70, 71, 83, 87 |
| 4 | 2 | 19610 | 0.9395 | 1.5969 | 0.9762 | 2, 5, 11, 12, 15, 23, 37, 43, 46, 53, 55, 60, 65, 72, 75, 77, 88, 90 |
| 4 | 3 | 20745 | 1.1558 | 2.0603 | 0.9550 | 9, 13, 16, 22, 24, 26, 36, 40, 45, 54, 56, 59, 64, 68, 69, 76, 80, 84 |
| 4 | 4 | 20320 | 1.2917 | 1.9986 | 0.9580 | 1, 4, 8, 10, 19, 27, 33, 39, 48, 49, 51, 57, 66, 67, 73, 78, 81, 89 |
| 4 | 5 | 20940 | 1.6708 | 2.7870 | 0.9252 | 6, 18, 20, 21, 25, 29, 31, 35, 38, 42, 44, 47, 63, 74, 79, 82, 85, 86 |
| 5 | 1 | 21000 | 1.1713 | 1.8911 | 0.9630 | 8, 11, 14, 18, 26, 30, 31, 33, 36, 40, 45, 52, 61, 68, 69, 72, 80, 84 |
| 5 | 2 | 20090 | 1.1360 | 1.9655 | 0.9603 | 1, 2, 15, 20, 23, 25, 34, 35, 44, 53, 58, 59, 70, 75, 76, 86, 88, 89 |
| 5 | 3 | 20330 | 1.1764 | 1.9768 | 0.9575 | 3, 5, 10, 13, 19, 28, 39, 42, 47, 49, 50, 51, 73, 74, 78, 81, 82, 85 |
| 5 | 4 | 20625 | 1.1907 | 1.9228 | 0.9664 | 7, 9, 17, 21, 22, 29, 38, 46, 54, 55, 56, 60, 62, 63, 66, 71, 77, 87 |

| Repeat | Fold | Test rows | MAE | RMSE | $R^2$ | Held out batch IDs |
|---|---|---|---|---|---|---|
| 5 | 5 | 20365 | 1.2481 | 2.0775 | 0.9579 | 4, 6, 12, 16, 24, 27, 32, 37, 41, 43, 48, 57, 64, 65, 67, 79, 83, 90 |

*MAE and RMSE are in g/L and pool the rows in each 18-batch fold. Every fold contains six batches from each normal operating regime. Fits within and across repetitions reuse the same 90 batches.*

**Table S4  Earlier prefix summary OOD warnings**

| Horizon h | Normal flagged | Normal rate | Deviation flagged | Deviation rate |
|---|---|---|---|---|
| 12 | 0/15 | 0.00% | 2/10 | 20.00% |
| 24 | 2/15 | 13.33% | 6/10 | 60.00% |
| 48 | 1/15 | 6.67% | 6/10 | 60.00% |
| 72 | 0/15 | 0.00% | 4/10 | 40.00% |

*Each horizon has a separate detector fitted to 75 normal development-batch summaries and a fitted-reference 95th-percentile threshold. These values are distinct from the final row-level OOD diagnostic.*

**Table S5 Earlier Random Forest empirical range evaluation**

| Fixed test set | Nominal target | Observed coverage | Mean full width g/L | Median width g/L | Mean tree SD |
|---|---|---|---|---|---|
| Normal Test | 90.00% | 84.89% | 5.034 | 3.744 | 1.069 |
| Deviation Test | 90.00% | 70.41% | 5.200 | 4.284 | 1.107 |

*The regressor was trained on 60 normal batches and scaling calibrated on 15 separate normal validation batches. Nominal coverage is not guaranteed for serially dependent rows or shifted deviation batches.*

**Table S6  Recorded fault reference windows**

| Batch | First activity h | Last activity h | Active episodes | Active rows | Batch rows |
|---|---|---|---|---|---|
| 91 | 20.0 | 214.0 | 3 | 203 | 1290 |
| 92 | 80.0 | 160.0 | 2 | 122 | 1150 |
| 93 | 70.0 | 90.0 | 1 | 101 | 1050 |
| 94 | 20.0 | 110.0 | 2 | 72 | 1150 |
| 95 | 20.0 | 211.0 | 3 | 188 | 1055 |
| 96 | 70.0 | 90.0 | 1 | 101 | 1150 |
| 97 | 20.0 | 214.0 | 3 | 203 | 1125 |
| 98 | 80.0 | 160.0 | 2 | 122 | 1150 |
| 99 | 20.0 | 110.0 | 2 | 72 | 1255 |
| 100 | 20.0 | 110.0 | 2 | 72 | 1150 |

*Activity means a non-zero recorded fault reference within a documented deviation batch. The first-to-last phase includes inactive gaps between episodes. Episode counts do not establish independent physical mechanisms.*

**Table S7 Numerical reproduction of the archived main run**

| Result category | Rows | Maximum absolute numeric difference |
|---|---|---|
| Per batch outer metrics | 200 | 0.000e+00 |
| Pooled outer metrics | 4 | 0.000e+00 |
| Outer held out predictions | 113,935 | 1.148e−41 |
| Final bundle diagnostics | 100 | 1.110e−16 |
| Held out risk summary | 4 | 0.000e+00 |
| Paired batch bootstrap | 2 | 0.000e+00 |

*The audit passed 239 checks and verified all five normal and ten deviation outer tests. Scientific cells were executed sequentially as Python after Jupyter-kernel startup was unavailable. This is numerical reproduction, not independent biological validation.*

**Table S8 Main run and software metadata**

| Item | Value | Interpretation |
|---|---|---|
| Main run mode | full | Reportable full run |
| Main regressor | HistGradientBoostingRegressor | 180 boosting iterations |
| Risk model | HistGradientBoostingClassifier | Uncalibrated warning score |
| Deviation weight multiplier | 3.0 | Not tuned in a nested outer loop |
| Normal outer folds | 5 | 18 complete normal batches per fold |
| Deviation outer folds | 10 | One documented deviation batch per fold |
| Main Python | 3.13.15 | Archived execution metadata |
| Main NumPy | 2.3.5 | Archived execution metadata |
| Main pandas | 2.2.3 | Archived execution metadata |
| Main scikit learn | 1.8.0 | Archived execution metadata |
| Earlier scikit learn | 1.6.1 | Baseline and diagnostic experiments |

*A smoke run is an installation check and is not the source of the reported result tables. Software versions should be retained with the frozen release.*

**Table S9  Final reliability diagnostics for deviation batches**

| Batch | Rows | Median risk | Warning rate | Median OOD | 95th percentile OOD | OOD flag rate |
|---|---|---|---|---|---|---|
| 91 | 1290 | 0.9771 | 91.86% | 0.4541 | 0.5847 | 3.02% |
| 92 | 1150 | 0.7856 | 65.22% | 0.4161 | 0.6686 | 11.83% |
| 93 | 1050 | 0.8557 | 70.76% | 0.4104 | 0.7041 | 9.05% |
| 94 | 1150 | 0.9491 | 89.39% | 0.4452 | 0.6841 | 12.17% |
| 95 | 1055 | 0.9741 | 90.05% | 0.4595 | 0.5767 | 2.27% |
| 96 | 1150 | 0.7588 | 67.22% | 0.3911 | 0.6959 | 7.83% |
| 97 | 1125 | 0.7957 | 89.78% | 0.3905 | 0.4927 | 1.96% |
| 98 | 1150 | 0.7033 | 63.74% | 0.3910 | 0.6377 | 6.26% |
| 99 | 1255 | 0.9098 | 89.72% | 0.4356 | 0.6358 | 5.98% |
| 100 | 1150 | 0.8390 | 86.17% | 0.4012 | 0.6201 | 4.61% |

*The OOD threshold was 0.626829. These final-bundle scores were calculated against the development reference and are deployment diagnostics, not independent estimates of detector accuracy.*

**Table S10  Initial selected Random Forest performance by test batch**

| Set | Batch | Rows | MAE | RMSE | $R^2$ |
|---|---|---|---|---|---|
| Deviation | 91 | 1290 | 5.7196 | 6.9945 | -7.0557 |
| Deviation | 92 | 1150 | 0.7016 | 1.1225 | 0.9821 |
| Deviation | 93 | 1050 | 1.0829 | 1.4991 | 0.9807 |
| Deviation | 94 | 1150 | 3.5769 | 4.8156 | -0.5033 |
| Deviation | 95 | 1055 | 3.1884 | 4.4600 | -0.5400 |
| Deviation | 96 | 1150 | 1.2855 | 2.3187 | 0.9421 |
| Deviation | 97 | 1125 | 1.1554 | 1.5579 | 0.9649 |
| Deviation | 98 | 1150 | 1.3007 | 2.1488 | 0.9651 |
| Deviation | 99 | 1255 | 3.0254 | 4.0526 | -0.0195 |
| Deviation | 100 | 1150 | 5.5517 | 7.5627 | -3.5315 |
| Normal | 2 | 1150 | 0.8120 | 1.1805 | 0.9878 |
| Normal | 5 | 895 | 0.3582 | 0.5235 | 0.9970 |
| Normal | 9 | 1260 | 3.4804 | 4.3381 | 0.1254 |
| Normal | 14 | 1150 | 0.8145 | 1.1910 | 0.9856 |
| Normal | 28 | 1150 | 1.1156 | 1.8933 | 0.9547 |
| Normal | 34 | 1150 | 1.1542 | 1.7728 | 0.9436 |
| Normal | 45 | 1310 | 3.4600 | 4.3842 | -0.1426 |
| Normal | 46 | 1150 | 0.5240 | 0.8084 | 0.9946 |
| Normal | 47 | 1140 | 1.4228 | 2.2373 | 0.9379 |
| Normal | 60 | 1150 | 1.5183 | 1.9939 | 0.9489 |
| Normal | 66 | 1150 | 0.9677 | 1.1972 | 0.9840 |
| Normal | 69 | 1125 | 0.5044 | 0.6514 | 0.9956 |
| Normal | 71 | 1070 | 1.1653 | 1.6866 | 0.9779 |

| Set | Batch | Rows | MAE | RMSE | $R^2$ |
|---|---|---|---|---|---|
| Normal | 77 | 1025 | 0.4919 | 0.7924 | 0.9932 |
| Normal | 83 | 1205 | 1.3354 | 1.8934 | 0.9593 |

*MAE and RMSE are in g/L. The selected Random Forest was fitted on the combined 75 normal development batches. These results precede the principal HGB comparison.*

**Table S11: Earlier Random Forest fault phase metrics by batch**

| Batch | Phase | Rows | MAE | RMSE | $R^2$ |
|---|---|---|---|---|---|
| 91 | Before onset | 99 | 0.0000 | 0.0000 | 0.9978 |
| 91 | First to last activity | 971 | 5.4201 | 6.5937 | -7.2169 |
| 91 | After last activity | 220 | 9.8314 | 9.9384 | -879.7705 |
| 92 | Before onset | 399 | 0.2609 | 0.3664 | 0.9865 |
| 92 | First to last activity | 401 | 0.4894 | 0.5644 | 0.9829 |
| 92 | After last activity | 350 | 1.6070 | 2.0826 | 0.3480 |
| 93 | Before onset | 349 | 0.1440 | 0.2021 | 0.9949 |
| 93 | First to last activity | 101 | 0.6874 | 0.7263 | 0.6958 |
| 93 | After last activity | 600 | 1.7402 | 1.9827 | 0.8610 |
| 94 | Before onset | 99 | 0.0000 | 0.0000 | 0.0970 |
| 94 | First to last activity | 451 | 0.4664 | 0.6485 | 0.9777 |
| 94 | After last activity | 600 | 6.4978 | 6.6459 | -12.7571 |
| 95 | Before onset | 99 | 0.0000 | 0.0000 | -0.1766 |
| 95 | First to last activity | 956 | 3.5149 | 4.6804 | -1.3355 |
| 96 | Before onset | 349 | 0.0555 | 0.0928 | 0.9989 |

| Batch | Phase | Rows | MAE | RMSE | $R^2$ |
|---|---|---|---|---|---|
| 96 | First to last activity | 101 | 0.5155 | 0.5934 | 0.7469 |
| 96 | After last activity | 700 | 1.9986 | 2.9634 | 0.5024 |
| 97 | Before onset | 99 | 0.0000 | 0.0000 | 0.8268 |
| 97 | First to last activity | 971 | 1.2154 | 1.5211 | 0.9629 |
| 97 | After last activity | 55 | 3.3518 | 3.7538 | -91.4119 |
| 98 | Before onset | 399 | 0.1505 | 0.2409 | 0.9955 |
| 98 | First to last activity | 401 | 1.4974 | 1.6383 | 0.8602 |
| 98 | After last activity | 350 | 2.2937 | 3.3027 | -1.3470 |
| 99 | Before onset | 99 | 0.0000 | 0.0000 | 0.7407 |
| 99 | First to last activity | 451 | 0.4762 | 0.7236 | 0.9748 |
| 99 | After last activity | 705 | 4.9807 | 5.2795 | -4.6009 |
| 100 | Before onset | 99 | 0.0000 | 0.0000 | -0.6381 |
| 100 | First to last activity | 451 | 0.5696 | 0.9424 | 0.9439 |
| 100 | After last activity | 600 | 10.1885 | 10.4191 | -42.0445 |

*These are held-out Random Forest results from the original fixed deviation stress test. The after phase is absent when no recorded observations occur after the last activity.*

**Table S12  Normal batch RMSE summaries by operating regime**

| Strategy | Regime | Batches | Mean RMSE | Median RMSE | Minimum | Maximum | Negative $R^2$ |
|---|---|---|---|---|---|---|---|
| Normal-only HGB | Recipe | 30 | 1.8844 | 1.6337 | 0.3288 | 4.7252 | 0 |
| Normal-only HGB | Operator | 30 | 2.0190 | 1.8606 | 0.5711 | 4.6407 | 1 |
| Normal-only HGB | APC | 30 | 1.0844 | 1.0943 | 0.3867 | 1.9357 | 0 |
| Fault-inclusive HGB | Recipe | 30 | 1.9615 | 1.8552 | 0.2906 | 4.5754 | 0 |
| Fault-inclusive HGB | Operator | 30 | 1.9570 | 1.7887 | 0.6072 | 4.5240 | 0 |
| Fault-inclusive HGB | APC | 30 | 1.1218 | 1.2048 | 0.3756 | 1.8320 | 0 |

*RMSE values are in g/L and are calculated separately for complete held-out normal batches. The full 180-row normal batch table is available in the machine-readable results.*

**Table S13  Earlier horizon specific OOD model audit**

| Horizon h | Summary inputs | PCA components | Explained variance | Fitted threshold |
|---|---|---|---|---|
| 12 | 105 | 14 | 95.819% | 0.516255 |
| 24 | 105 | 15 | 95.929% | 0.526419 |
| 48 | 105 | 16 | 95.913% | 0.510930 |
| 72 | 105 | 15 | 95.312% | 0.566229 |

*A separate preprocessing and Isolation Forest pipeline was fitted at each horizon using the 75 normal development batches.*

**Table S14 Additional prediction and reliability diagnostics**

| Diagnostic | Count | Value or interpretation |
|---|---|---|
| Normal-only held-out predictions below zero | 686 | -0.3164 g/L |
| Fault-inclusive held-out predictions below zero | 710 | -0.2640 g/L |
| Normal empirical absolute-error radius | Not applicable | 3.0398 g/L |
| Deviation empirical absolute-error radius | Not applicable | 3.1164 g/L |
| Final OOD reference rows | 22500 | Maximum 250 rows per normal batch |
| Final OOD threshold | Not applicable | 0.626829 |

*Negative predictions were retained for all reported error metrics. Empirical radii are descriptive and are not formal confidence or safety limits.*

**Table S15 Machine-readable result inventory**

| Directory | File | Purpose |
|---|---|---|
| Baseline | validation_results.csv | Validation metrics for dummy, Linear Regression and Random Forest |
| Baseline | normal_and_fault_test_results.csv | Selected Random Forest pooled test metrics |
| Baseline | normal_test_predictions.csv | Held-out normal row predictions |
| Baseline | fault_test_predictions.csv | Held-out deviation row predictions and reference fields |
| Baseline | normal_test_metrics_by_batch.csv | Normal test metrics by batch |
| Baseline | fault_test_metrics_by_batch.csv | Deviation test metrics by batch |
| Experiments | experiment_1_repeated_batch_cv_fold_metrics.csv | All 25 repeated complete-batch folds |
| Experiments | experiment_2_time_feed_ablation.csv | Pooled ablation metrics |
| Experiments | experiment_3_fault_phase_overall.csv | Pooled and mean-batch phase summaries |
| Experiments | experiment_3_fault_phase_by_batch.csv | Phase metrics for every available batch phase |
| Experiments | experiment_3_fault_onset_audit.csv | Recorded first and last activity and episode counts |
| Experiments | experiment_4_model_comparison_test.csv | Fixed-test model-family metrics |
| Experiments | experiment_5_early_ood_scores.csv | All horizon and batch OOD scores |

| Directory | File | Purpose |
|---|---|---|
| Experiments | experiment_5_ood_evaluation_summary.csv | Horizon-specific warning counts |
| Experiments | experiment_5_uncertainty_summary.csv | Earlier empirical-range coverage summary |
| Main | cross_validated_predictions.csv | All 113,935 outer-held-out predictions and risk scores |
| Main | cross_validated_overall_metrics.csv | Principal pooled metrics |
| Main | cross_validated_batch_metrics.csv | Principal metrics for both models and all batches |
| Main | paired_batch_bootstrap.csv | Paired batch RMSE summaries |
| Main | fault_risk_summary.csv | Held-out risk warning summaries and within-fault AUC |
| Main | deployment_reliability_by_batch.csv | Final-bundle risk and OOD diagnostics by batch |
| Main | run_metadata.json | Features, settings, weights versions and diagnostic thresholds |

*These files provide the numerical basis of the manuscript. Plot files and human-readable README guides accompany them in the repository. The original data source remains the cited Mendeley Data record [6].*

**Supplementary Figures**

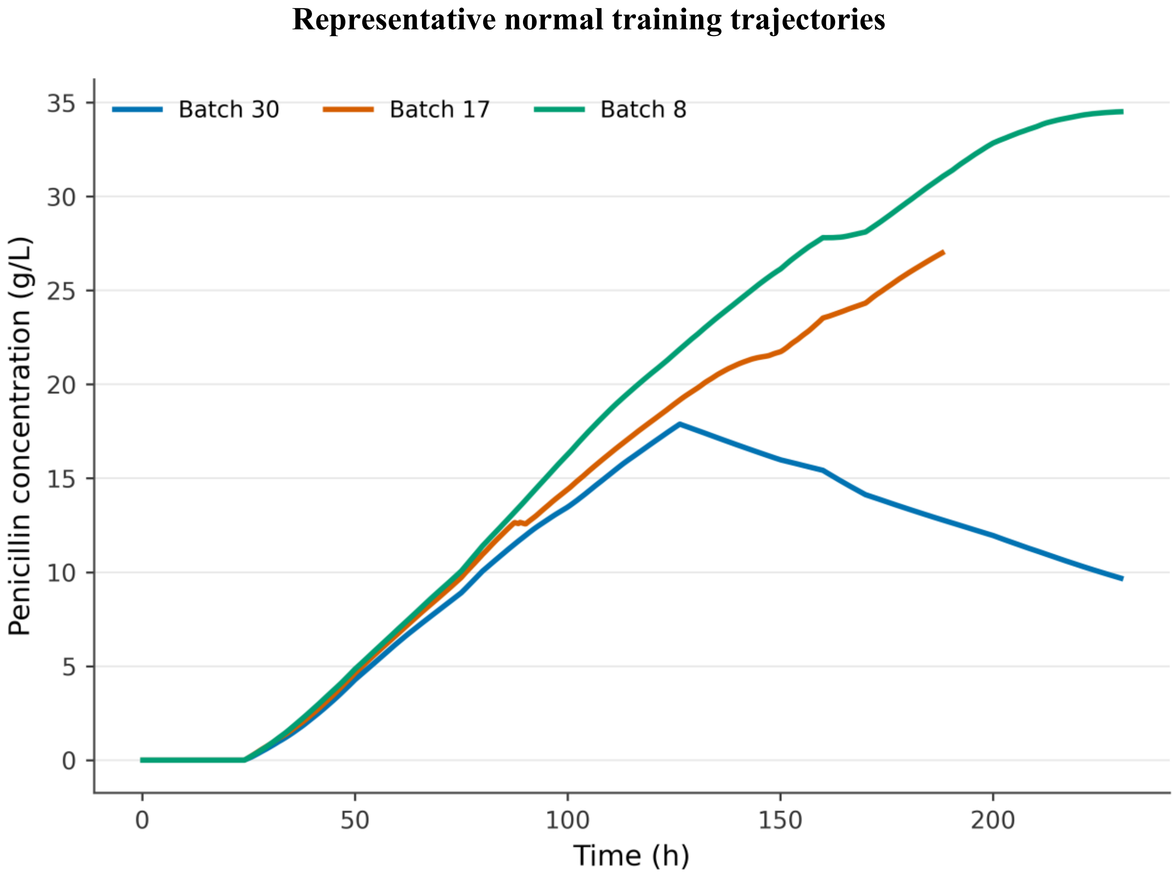


**Figure S1:** Penicillin concentration trajectories for baseline training batches 30, 17 and 8, the three batches used in the original visual inspection. They illustrate different normal trajectory lengths and shapes. The panels are descriptive and were not selected as independent evidence of model accuracy.

### Initial Random Forest actual versus predicted concentrations

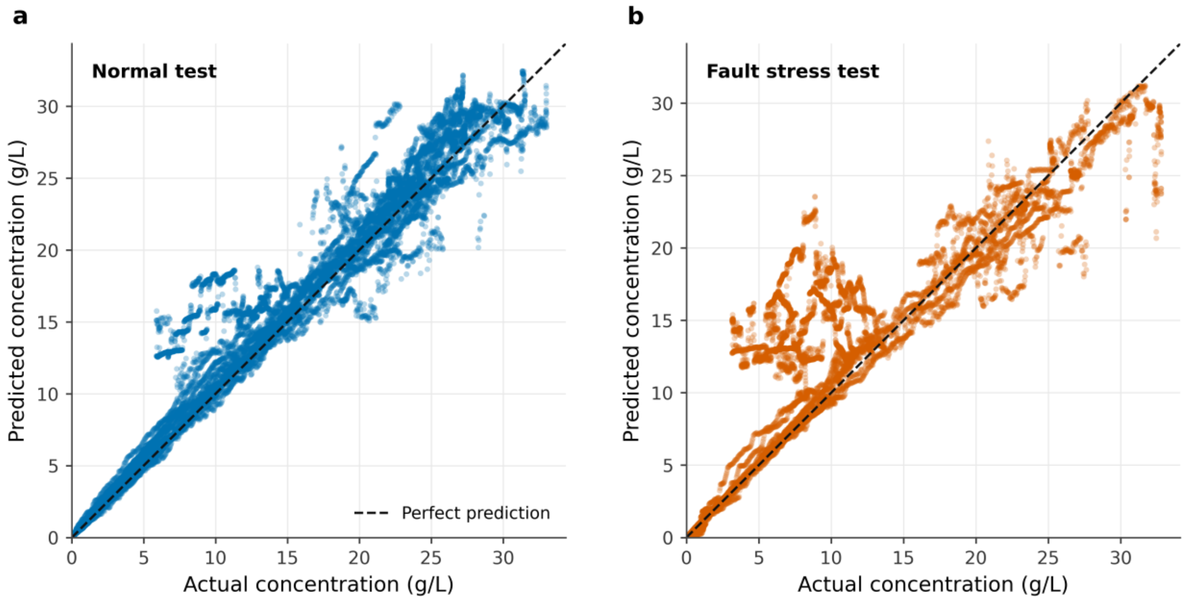


**Figure S2:** Actual-versus-predicted concentration for the initial selected Random Forest on the fixed 15-batch normal test set and ten-batch deviation stress test. Dashed lines denote equality. Points are time observations nested within batches. The normal-trained model shows visibly larger dispersion and systematic error on deviation trajectories.

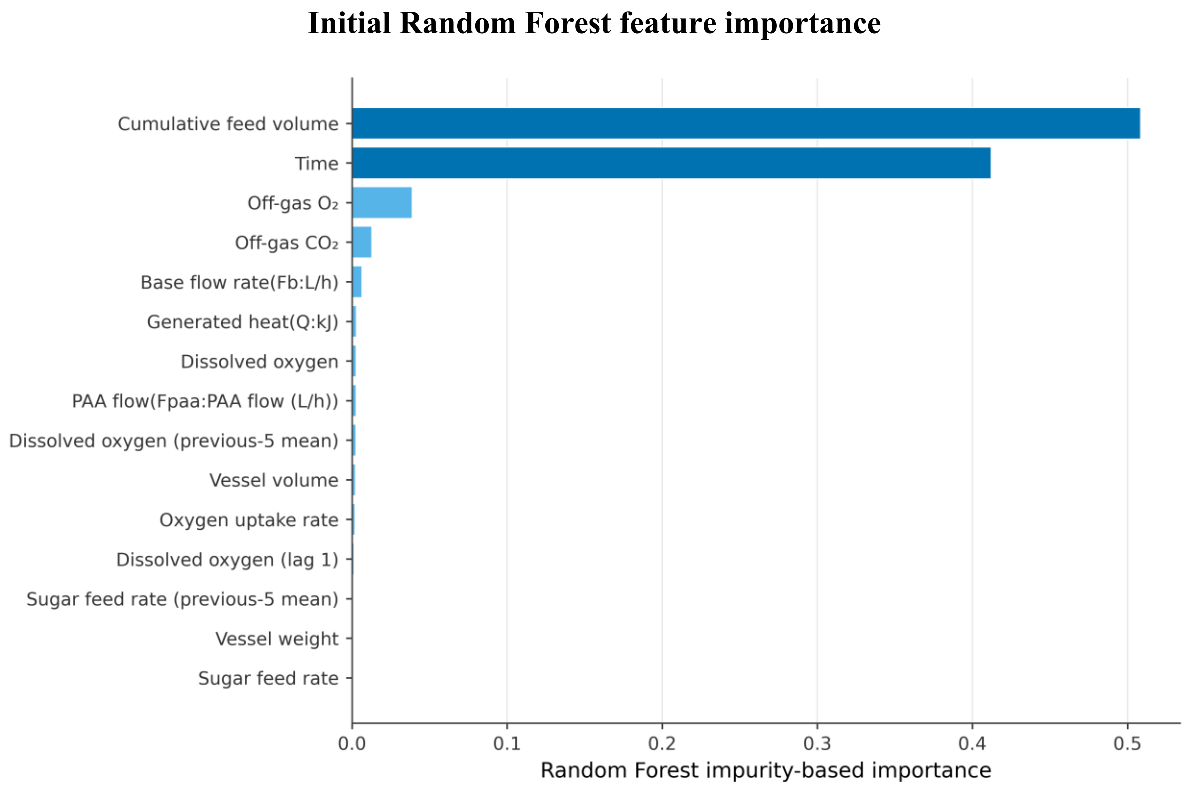


**Figure S3:** Top 15 impurity-based feature importances from the selected Random Forest. Cumulative sugar feed and elapsed time account for 92.06% of the total fitted importance. The chart describes the fitted model and does not establish causal biological importance; correlated variables can redistribute importance.

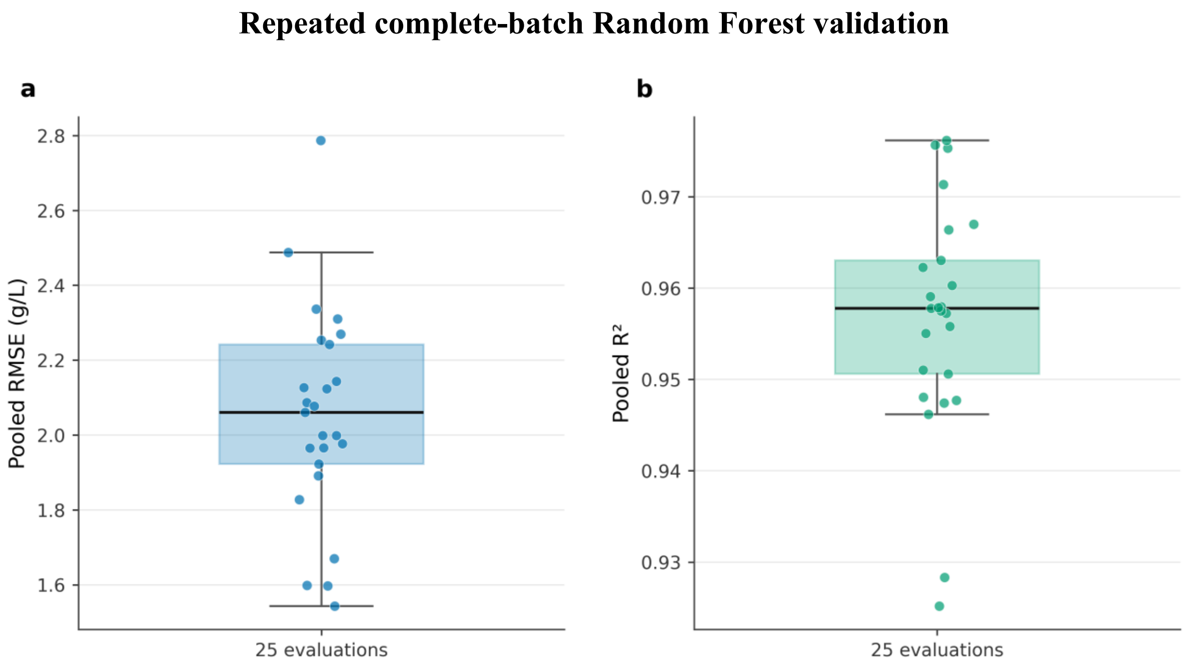


**Figure S4:** Boxplots and overlaid fold points for pooled RMSE and $R^2$ across five repetitions of five regime-stratified complete-batch folds. Each fit trains on 72 normal batches and tests 18. Mean RMSE is 2.050 g/L, standard deviation 0.286 g/L and range 1.543–2.787 g/L. The 25 folds are dependent because they reuse the same 90 batches.

#### Time and cumulative feed ablation

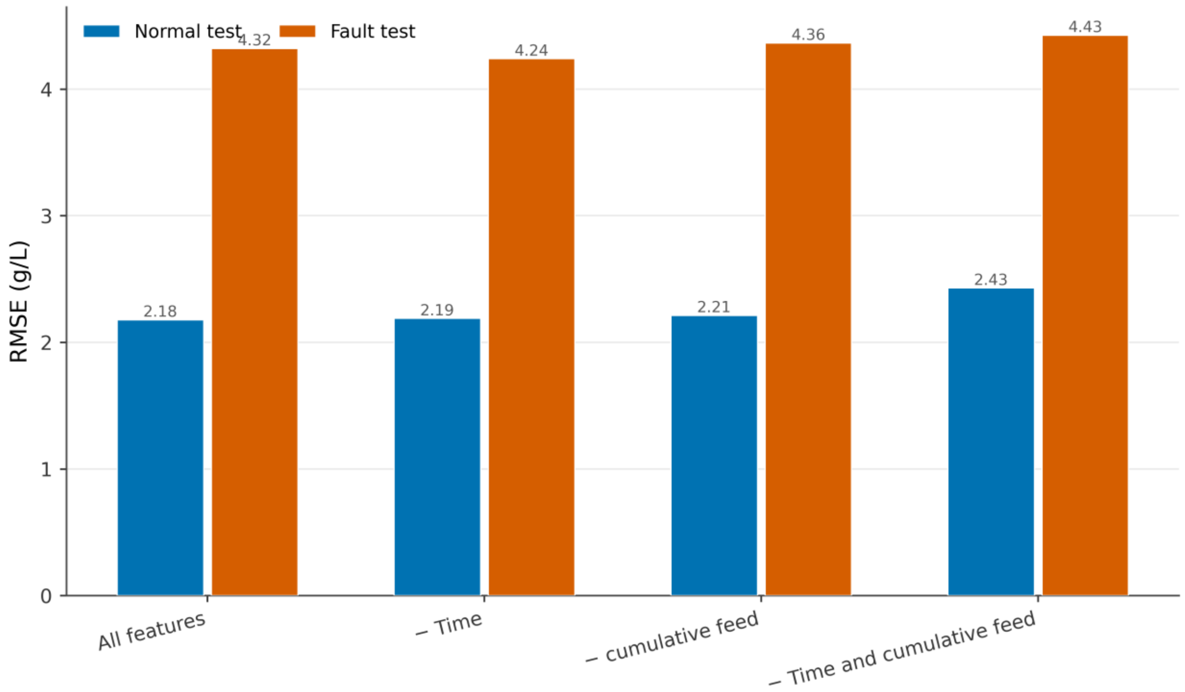


**Figure S5:** Earlier Random Forest ablation using all inputs, without time, without cumulative feed and without both. Values are pooled RMSE on the fixed normal and deviation test sets. Removing both increases normal RMSE by 11.53% and deviation RMSE by 2.44% relative to this experiment's all-input fit. Other progress proxies remain available.

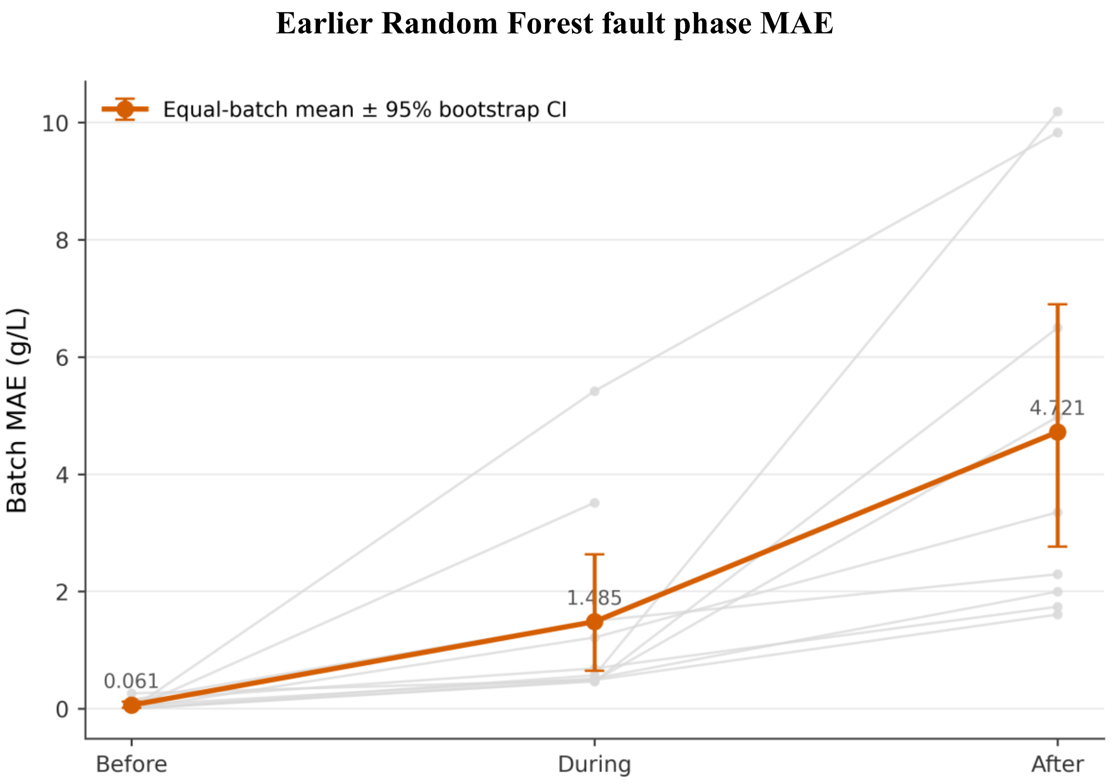


**Figure S6:** Grey lines connect individual deviation batches, and the red series shows mean batch MAE with the earlier notebook's batch-bootstrap interval. Mean batch MAE is 0.061, 1.485 and 4.721 g/L before, within and after the first-to-last recorded activity window. The after phase contains nine batches. Phase differences are descriptive and confounded with time and target range.

**Earlier, a normally trained model comparison**

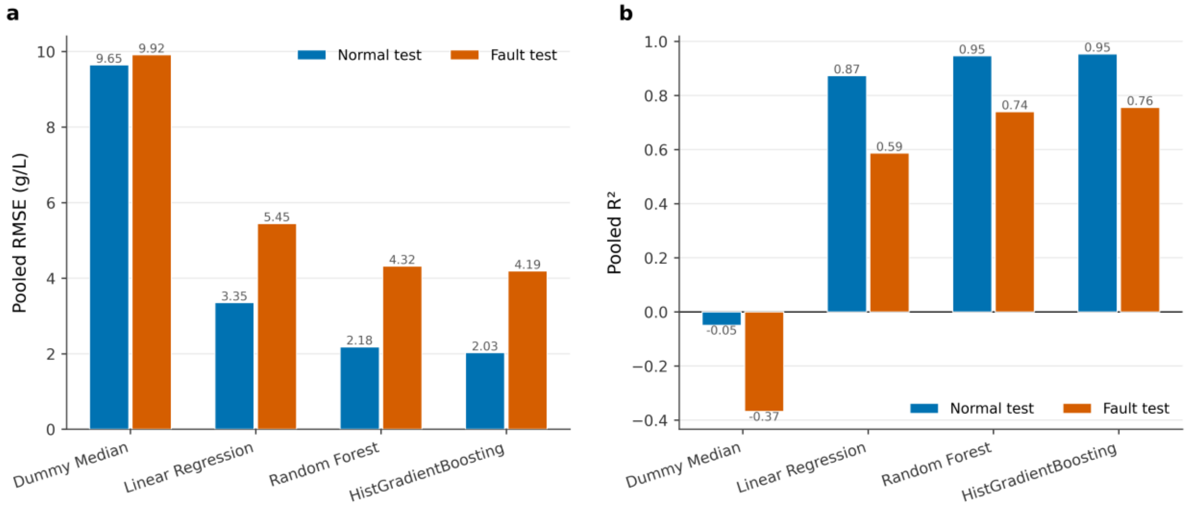


**Figure S7:** Pooled RMSE and $R^2$ for the median dummy, Linear Regression, Random Forest and HGB on fixed normal and deviation test batches. HGB uses 300 iterations here and is not the final HGB configuration. Every learned model performs worse than normal; differences from the principal analysis cannot be attributed to a single component because the training and validation designs differ.

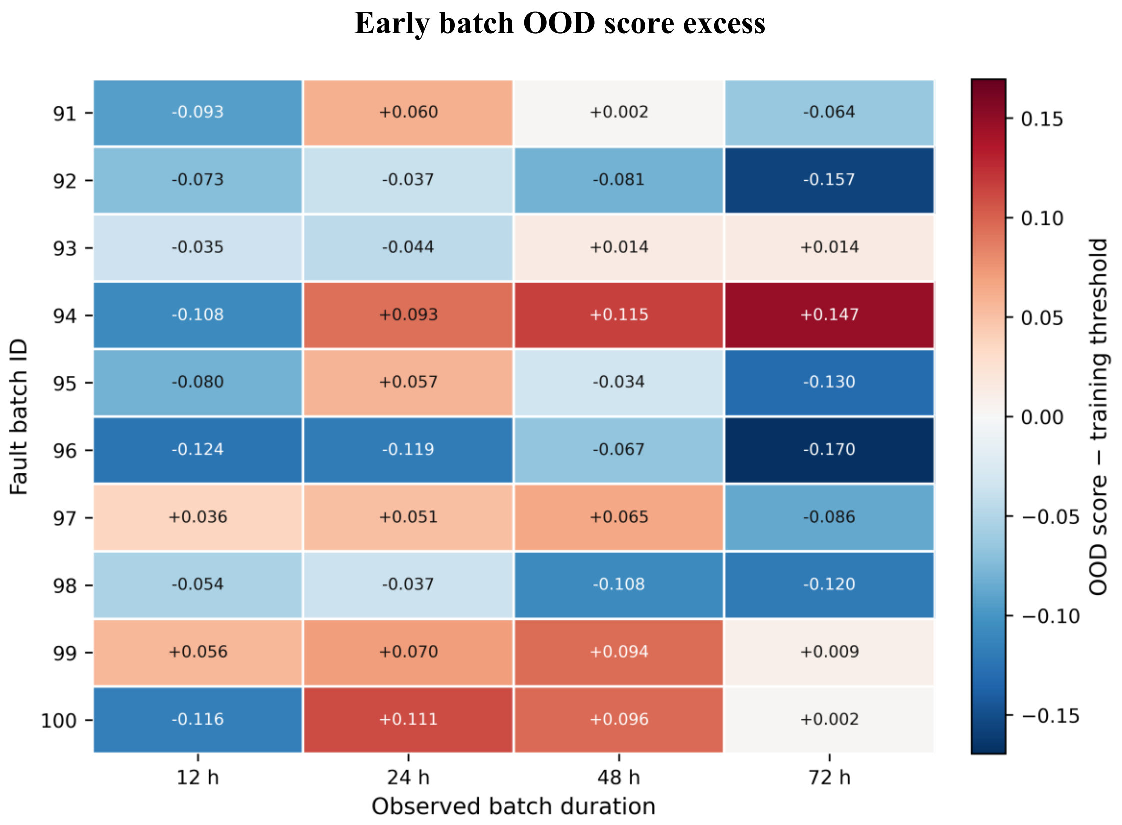


**Figure S8:** Horizon-specific OOD score minus its fitted 95th-percentile normal threshold for batches 91–100 at 12, 24, 48 and 72 h. Positive cells generate an OOD warning. Each detector uses 75 normal development-batch summaries. A warning denotes an unusual prefix summary, not a confirmed process fault or concentration-error forecast.
